# The identification and analysis of meristematic mutations within the apple tree that developed the *RubyMac* sport mutation

**DOI:** 10.1101/2023.01.10.523380

**Authors:** Hequan Sun, Patrick Abeli, José Antonio Campoy, Thea Rütjes, Kristin Krause, Wen-Biao Jiao, Maria von Korff, Randy Beaudry, Korbinian Schneeberger

## Abstract

Understanding the molecular basis of sport mutations in fruit trees can accelerate breeding of novel cultivars. For this, we analyzed the DNA of the apple tree that evolved the *RubyMac* phenotype through a sport mutation that introduced changes in fruit coloration in upper branches of the tree. Unexpectedly, we not only found 46 *de novo* mutations, but also 54 somatic gene conversions (i.e., loss-of-heterozygosity mutations) distinguishing the mutant and wild-type branches of the tree. Approximately 30% of the *de novo* mutations and 80% of the gene conversions were observed only in specific cells layers suggesting that they occurred in the corresponding meristematic layers. Interestingly, the *de novo* mutations were enriched for GC=>AT transitions, while the gene conversions showed the opposite bias for AT=>GC transitions suggesting that GC-biased gene conversions have the potential to counteract the AT-bias of *de novo* mutations. By comparing the gene expression patterns in fruit skins from mutant and wild-type branches, we found 56 differentially expressed genes including 18 that were involved in anthocyanin biosynthesis. While none of the differently expressed genes harbored a mutation, we found that some of the mutations affected the integrity of candidate genes in regions of the genome that were recently associated with natural variation in fruit coloration.

## INTRODUCTION

In plants, a somatic mutation introduces a change in the DNA of an individual, somatic cell and, in consequence, all cells that are derived from it. If a somatic mutation occurs in the meristem, the mutation can be propagated into large sectors of the plants. However, as the identities of the meristematic layers remain mostly intact during organ development (i.e., the different cell layers of an organ usually develop from of different meristematic layers), some of the somatic mutations in the meristem (meristematic mutations) might be specific to certain cell layers despite being present in all cells of a newly develop organ.

Somatic mutations have been highly valuable in fruit tree breeding, as they can generate or improve agronomically important traits, and if observed in elite material, they do not even need to be introgressed to generate new cultivars. Bud sports in fruit trees are usually clonally propagated which keeps their somatic genomes intact and any derived somatic mutation can therefore also be passed on the next clonal generation^7–13^.

Examples for bud sports include changes in fruit coloration, which is not only an important trait to meet consumers’ preference but might also relate to chemical compounds benefiting human health^14–19^. Changes in the fruit coloration usually result from changes in the accumulation of anthocyanins, which are synthesized by enzymes, which are regulated by *MYB* transcription factors^20–24^ which could and have been changed by somatic mutations. Other examples of bud sport mutation in fruit trees as well as other plants have been recently reviewed by Foster and Aranzana^8^.

But despite their economic importance, the mutational mechanisms that lead to bud sport mutations is still far from being understood. Here, we identified and analyzed the somatic mutations of an apple tree (*Malus domestica* cultivar *McIntosh RubyMac*) that develops fruits with dark red skin in its upper (mutant) scaffolds, while the apples in the lower (wildtype) scaffolds remain pale red. Comparing 12 whole-genome sequencing datasets generated from three different types of tissues from four different regions of the tree, we found 100 somatic mutations including 53 that separated the mutant and wild-type branches of the tree. Unexpectedly, the mutations did not only introduce novel variation through spontaneous mutations (46), but included a similar number of loss-of-heterozygosity mutations (54), where wild-type heterozygous sites change to homozygous sites. The two types of mutations show opposing spectra and could be observed in different tissues suggesting that different types of mutational mechanisms act during the generation of somatic variation. Additional analysis of differentially expressed genes between the mutant and wildtype fruit peel revealed 56 genes with differential expression profiles including 18 involved in flavonoid or anthocyanin biosynthesis and regulation. While none of the differentially expressed genes was an obvious target of the somatic mutations, we identified mutations in other genes, including some in regions of the genome that were recently associated with natural variation in skin coloration.

## RESULTS

### Identification of meristematic mutations

The apple tree that evolved the *RubyMac* phenotype is growing in Michigan, USA (43°04’53.1"N 85°43’13.5"W). The top seven scaffolds show purple pigmentation on stamens and dark red fruits, which cannot be observed on the lower branches of the tree, where stamens remain white and fruits develop skin with pale red color (Fig. 1; Supplementary Figure 1). The acquired phenotype is stable even after clonal propagation, and reversions have been seldomly reported. This suggested that a genetic mutation could be responsible for the change in fruit coloration. As this change was present in the entire upper part of the tree including unconnected scaffolds, we assumed that this mutation occurred within the shoot apical meristem during the development of the stem and thereby separated the lower (wildtype) and upper (mutant) parts (Fig. 1a).

**Figure 1.**
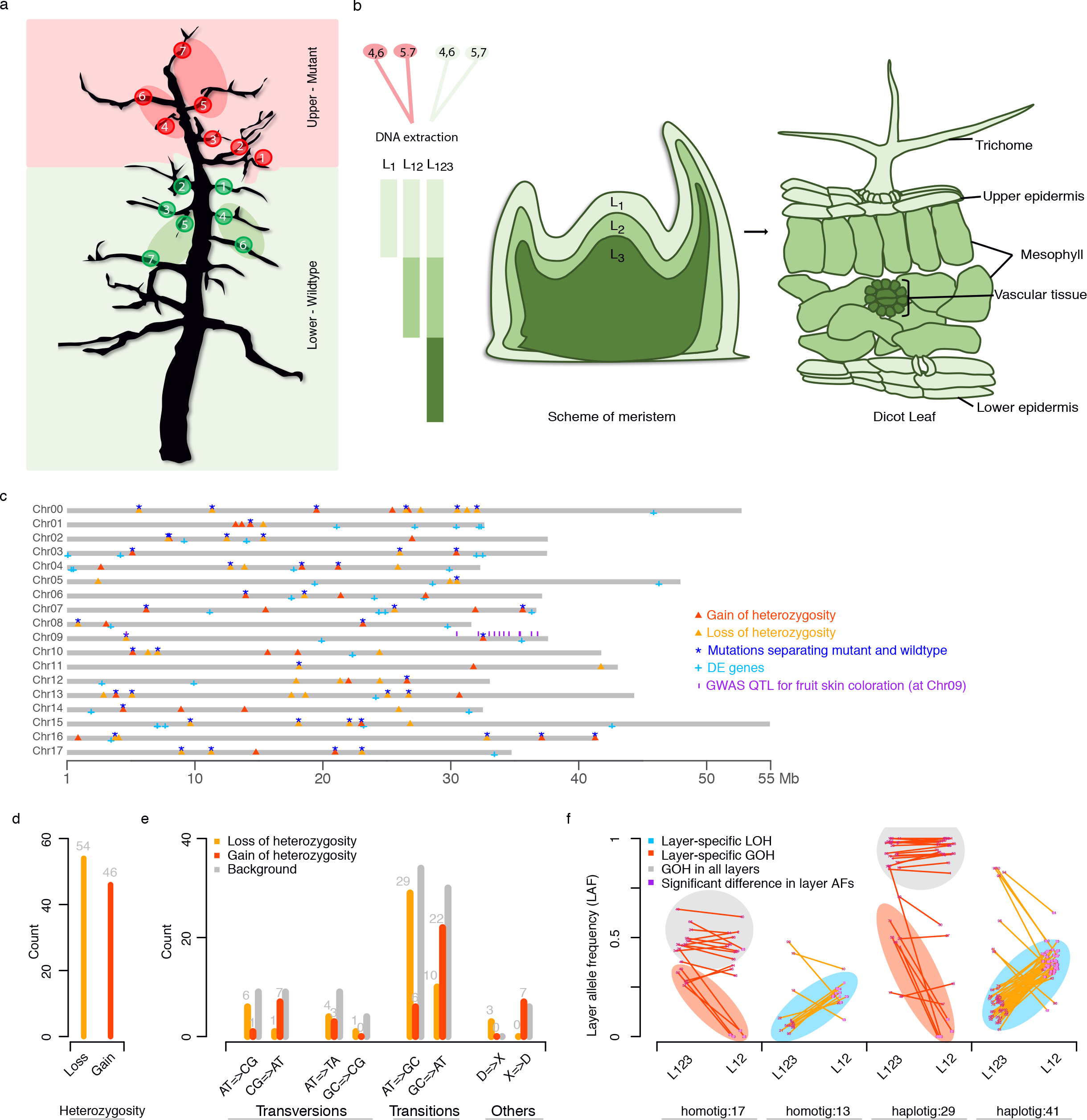
Schematics of the layer-specific whole genome sequencing based on a 3-layer meristem model, and characterization of somatic mutations. ***a***.Two-dimensional projection of the tree showing the sampling points for whole-genome as well as PCR amplicon sequencing. The pink background marks the scaffolds developing red apples, while the green background marks the scaffolds with pale red apples. The red and green eclipses show the approximated regions where the bulked leaves were selected for whole-genome sequencing. Mutant branches 1-7, wildtype branches 1-6, pooled mutant branches 4,6 and 5,7 (leaf, petiole), wildtype branches 4,6 and 5,7 (leaf, petiole) were genotyped with PCR amplicon sequencing. ***b***. Meristem model with three different layers and how the layers correspond to the cell layer of a leaf, i.e., *L_1_*: trichome and epidermis, *L_2_*: Mesophyll and *L_3_*: vascular tissue. ***c***.Distribution of 86 somatic mutations across the GDDH13 v1.1 genome^40^ (the sequence of the remaining 14 mutations could not be reliably mapped to the reference sequence). ***d***. 54 loss-of-heterozygosity (LOH) and 46 gain-of heterozygosity (GOH) were observed. ***e***. Mutational spectra of LOH and GOH mutations. The background distribution is based on germline variation (SNPs) found between both haplotypes of the apple tree. ***f***. LOH and GOH occurred in three major clusters in respect to the read frequency within two different types of samples (*L_123_* and *L_12_* samples) indicating their occurrence in different cell layers (Material and Methods). The clusters were consistently found on *haplotigs* as well as *homotigs* (where the reads of the other allele were aligned as well). The grey cluster shows *de novo* mutations that occur in almost all reads of the haplotype independent of the actual sample (*L_123_* or *L_12_*). The red cluster shows *de novo* mutations that occurred in approximately half of the reads in the *L_123_* samples but were almost entirely absent in the *L_12_* samples. The blue cluster shows loss-of-heterozygosity mutations which were observed at low frequency in *L_123_* and at intermediate frequency in the *L_12_* samples (which is indicative for mutations that occurred in *L_1_* or *L_2_*).

To identify meristematic mutations that occurred during the development of the upper mutant part, we sampled four different sets of leaves: leaves pooled from connected scaffolds of the mutant part of the tree (2x; where the scaffolds of both sets were distinct) as well as leaves pooled from connected scaffolds from the lower wildtype part of the tree (2x; where the scaffolds of both sets were again distinct) (Fig 1a). Pooling the DNA of the leaves of each set diluted the signal of somatic mutations that occurred in a few differentiated cells (e.g. within a single leaf), while mutations that occurred in the meristem and that are propagated to the entire upper tree would be present in both samples of the upper part. We therefore assumed that the mutations that we identified in both samples of upper part of the tree emerged in the meristems during stem development and that those could be candidate mutations for the sport mutation phenotype.

Following a meristematic model that supports the presence of three layers^2^, we extracted DNA from three different leaf sub-tissues (of each of the four sampled leaf sets) to enrich for cells that originated from different meristematic layers leading to a total of 12 DNA samples (Fig. 1b). The different extractions included DNA from trichomes (enriched for cells derived from layer 1 (or *L_1_*)), peeled leaf surface (enriched for cells derived from layer 1 and 2 (or *L_12_*)) and whole leaf blades (including cells derived from all three layers (or *L_123_*)) (Fig. 1b). We sequenced the *L_123_* and *L_12_* samples with 69-87x genome coverage using Illumina 2×250 bp short-reads (Materials and Methods). The sequencing of the four trichome *L_1_* DNA samples yielded only 4-7x of the apple genome as there was substantial pathogen contamination in the sequencing data (Supplementary Table S1).

Using the whole genome sequencing data of one of the wildtype samples, we generated a *de novo* assembly of the *RubyMac* genome using *DiscovarDeNovo*^34^ with a total length of 875 Mb and a contig *N50* of 18.8 kb (referred to as *RubyMac DDN* assembly). The assembly size was larger than the haploid genome size of 731 Mb as estimated by *findGSE*^41^ and also larger than the reference assemblies of *Golden Delicious*^39–40^. We speculated that this was due to the high heterozygosity of *RubyMac* (1.2±0.1% estimated by *findGSE*), resulting in many haplotype-specific contigs (or *haplotigs*). Using a computational pipeline to identify haplotigs^42^, we characterized haplotigs with a combined length of ~242 Mb within the *RubyMac DDN* assembly explaining the inflated assembly size (Materials and Methods). We also extracted RNA from peels of both mutant and wildtype fruits (each with three replicates), from which 24-27 million Illumina 150 bp single-end reads were respectively sequenced. With the guidance of RNA-seq data, we predicted 42,981 high confidence gene models in the *RubyMac DDN* assembly using *augustus*^44,45^ (Materials and Methods).

We aligned each of the 12 sequencing read sets to the *RubyMac DDN* genome with an alignment rate of 92-96%, which was much higher than with any other apple reference sequence, suggesting that this was the optimal reference sequence for mutation identification in our data despite that the assembly was not on chromosome level (Supplementary Table S1).

By comparing the eight *L_12_* and *L_123_* samples, we defined an initial set of 69 candidate mutations, which consisted of three subsets, which were defined based on three different quality metrices. We performed Sanger sequencing with DNA sampled from single leaves taken from seven upper and six lower scaffolds of the tree for each mutation to test the effect of each of the quality metrics on the success of the mutation identification (Material and Methods; Supplementary Figure 2-30). The first set consisted of 29 mutations, which were called in more than two samples with an alternative base quality above 30 in at least two of the samples. Of those we could confirm 27 mutations. The second set consisted of 24 mutations, which were also predicted in more than two samples but with less than two samples with an alternative base quality above 30. Of those, only two were confirmed by Sanger sequencing. The third, least stringent, set included six mutations, which were predicted in only up to two samples with at least one sample with an alternative base quality above 30. None of them could be confirmed by Sanger sequencing. For the remaining 10 cases the PCR sequencing did not yield conclusive results.

**Figure 2.**
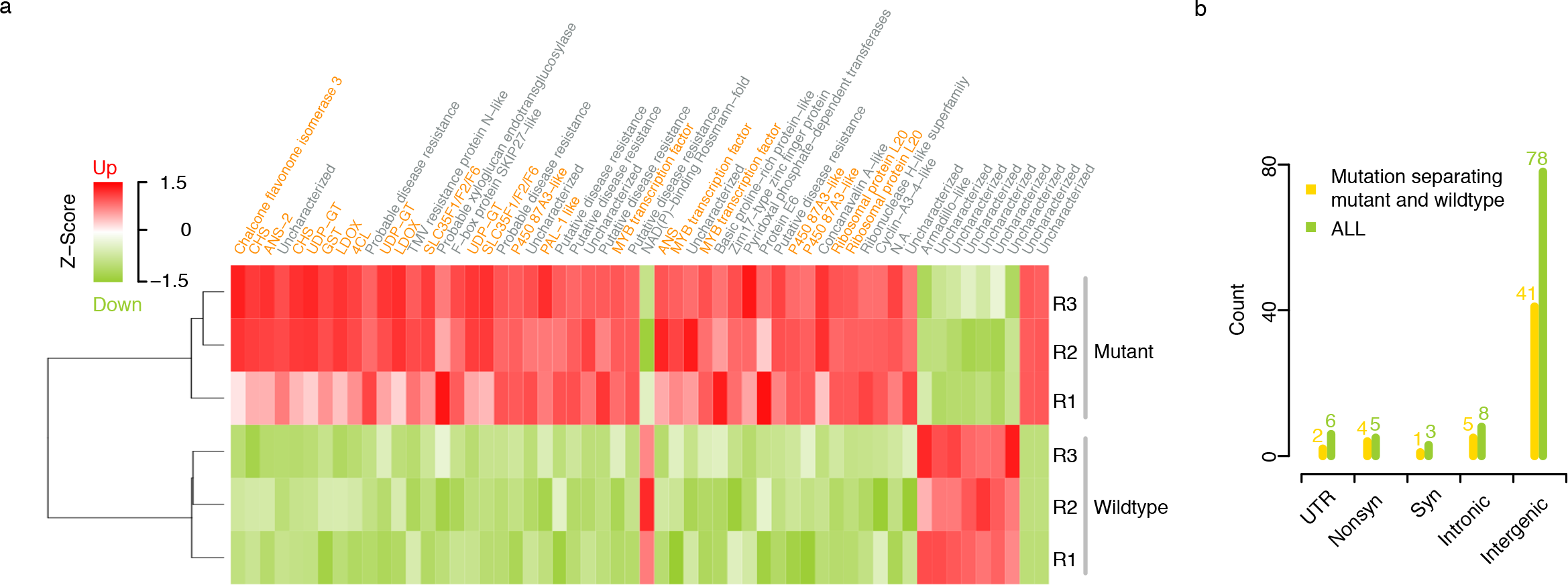
Differential gene expression analysis between mutant and wild-type fruit peel. *a*.56 differentially expressed genes (DEGs) including 48 genes up-regulated and 8 genes down-regulated in the mutant peel. Genes labeled with orange names were involved in flavonoid or anthocyanin biosynthesis and regulation (including MYB transcription factors). ***b***. Distribution pattern of mutational effects on gene integrity.

While it was obvious that the criteria for the second set were not sufficient to overcome a high rate of false positive, we found that among the eight unconfirmed cases in the first and third subset were five low-frequency mutations, which potentially could have even been missed with Sanger sequencing. For example, one of those mutations (a loss-of-heterozygosity mutation) was found with a minor allele frequency of less than 0.1 in the whole-genome sequencing data (Supplementary Figure 31) and was not confirmed by Sanger sequencing. However, within RNA-seq data that we generated from the peels of fruits, all three wildtype replicates again showed 4~6% reads (read depths: 595-784x) with the alternate allele, while not a single such read showed up in any of the three mutant replicates (read depths: 660-861x; Materials and Methods; Supplementary Figure 31). The identification of the alternate allele in RNA-seq suggested that this was a real difference between the mutant and wildtype branches despite it was missed in the Sanger sequencing.

Taken together, the high false positive rate in second set indicated that mutations with comparably low alternative base quality can usually not be trusted in our dataset. Among the other mutations in other two subsets, we were able to confirm 28 cases with either Sanger or RNA-seq data suggesting the minimum requirement on the base quality that we need to apply to guarantee satisfactory accuracy in predicting mutations, while some of the low-frequency candidate mutations might have been missed in the Sanger sequencing. Therefore, by applying these base quality requirements of the first and third set, we performed a second round of mutation identification where we identified 100 single-nucleotide mutations, which were all checked with visual inspections of their alignments. For further confirmation we also called mutations with the reference-based tool *MuTect*^36^ and the reference-free tool *discosnp*++^46^ and found all 100 mutations with both tools (Materials and Methods). Note, the *L_1_* samples were generally not used for mutation identification as the read coverage was too low for finding new mutations.

Among the 100 mutations, 53 mutations were common to both samples of the upper or common to both samples of the lower parts of the tree (and therefore separated the mutant and wild-type parts, while the remaining 47 mutations were not specific to the lower or upper parts, but occurred only in one region of the tree (10 mutations), or showed a cryptic distribution across mutant and wild-type (37 mutations; Supplementary Table S2). Using the chromosome-level reference assembly of apple^40^ as a guide, we found that the mutations were scattered across all chromosomes (Fig. 1c).

Besides small-scale nucleotide-level mutations, we also searched for structural variations (SV), and the movement of transposable elements (TE; Materials and Methods). However, despite an extensive search using SV detection tool like *CNVnator*^60^ as well as customized analysis of the depth of short-read alignments across samples followed by manual checking using alignment visualizations with IGV, we could not identify a single large-scale SVs or TE movement that would have distinguished the mutant and the wildtype branches of the tree.

### GC-biased gene conversions counteract AT-biased de novo mutations

*De novo* mutations (DNM) introduce novel variation, which initially is specific to one of the two homologous chromosomes (here referred to as gain-of-heterozygosity mutations). In rare cases DNM can occur at wildtype heterozygous sites and turn a heterozygous into a homozygous site (including either reversion to the wildtype allele or fixation of the derived allele, referred to as loss-of-heterozygosity mutations) or turn a heterozygous allele into a different heterozygous allele (referred to as change-of-heterozygous-allele mutations). However, the probability for loss-of-heterozygosity or change-of-heterozygous-allele mutations to occur due to spontaneous changes is very low as heterozygosity typically includes only a few percent of the nucleotides in a genome.

We defined a mutation as a gain-of-heterozygosity mutation if it was covered by more than 15 reads in at least one sample (which could be either mutant or wildtype) and at least three wildtype samples did not support the alternative allele with more than two reads. In contrast, a mutation was defined as loss-of-heterozygosity if the alternative allele was found in at least three wildtype samples with a coverage of over 15 while all four mutant samples did not feature more than 3x of the mutant allele. With these strict definitions all 100 mutations could be assigned as either loss or gain-of-heterozygosity (Supplementary Table S2).

Surprisingly, in contrast to what was expected, we observed a striking overrepresentation of loss-of-heterozygosity mutations. Overall, among the 100 mutations, we found 54 loss-of-heterozygosity and 46 gain-of-heterozygosity (Fig. 1c-d), even though the genome-wide heterozygosity was as low as 1.2%. It is important to note that all mutations were identified with the same computational procedure (comparing samples against other samples – independent of whether they are of mutant or wild-type origin), and there is no reason to expect differences in the detection performance between the different types of mutations.

But despite the unexpected overrepresentation of loss-of-heterozygosity mutations, change-of-heterozygous-allele mutations were completely absent. This suggested that it was actually not DNM that introduced the loss-of-heterozygosity mutations, but that they were introduced through the replacement of one allele by its homologous allele through gene conversions.

To understand more about the differences between DNM and gene conversions, we analyzed their mutational spectra (Fig. 1e). In general, DNM were highly enriched for GC=>AT transitions (48%; 22 out of 46) which recapitulated the spectrum of *de novo* mutations that were inherited through the germline^25^. In contrast, 54% (29 out of 54) of the gene conversions were AT=>GC transitions, while all the other types of gene conversions were underrepresented (2-19%). Interestingly, the contrasting spectra of genomic mutations and gene conversions, i.e., GC-biased gene conversions and AT-biased mutations, suggested a mechanism that could stabilize the GC content of a genome.

### Layers-specific de novo mutations and gene conversions

Meristematic mutations can be propagated into large parts of a plant and thereby introduce mutant sectors^47^. However, as the meristem is organized in layers^2^, meristematic mutations could be propagated into specific cell lineages of the sectors that they can be found in. In turn, this implies that somatic mutations could only be present in a fraction of the cells despite being present in large areas of the tree. The actual mutant allele frequency in the whole-genome sequencing data therefore depends on the fraction of cells of the mutated layer in the sequenced sample. In addition, the mutant allele frequency in the aligned reads also depends on the reference sequence, which either allows the alignment of the reads of both alleles (in the case the reference sequence is assembled as *homotig*) or which only allows the alignments of only one of the alleles (in the case the reference sequence is assembled as *haplotig*) in the respective region.

The estimation of the mutant frequencies is generally more accurate on *haplotigs* as the reads of the alleles derived from the homologous chromosome do not dilute the signal as compared to the estimation made with the alignments to *homotigs*. Among the 46 DNM, 29 were found on haplotigs suggesting that all the aligned reads were only derived from one allele. Among those, 9 (31%) DNM could be observed in only some of the reads suggesting that they were layer-specific (in fact, five of these layer-specific mutations could even be assigned to an actual layer as they were supported by half of the reads in the *L_123_* samples, while the mutant alleles were virtually absent in the *L_12_* samples suggesting that they occurred in *L_3_*) (Fig. 1f). In contrast, the remaining 20 (69%) DNM could be observed in nearly all the reads in both the *L_123_* and the *L_12_* samples suggesting that these mutations were in fact not layer-specific, but that they were present in all layers. It has previously been reported that meristematic cells can migrate between layers and that layer identity is not absolute^8^. The mutant allele frequencies of the 17 DNM detected on *homotigs* revealed the same types of mutations (with the difference that the mutant allele frequencies were generally only half of the allele frequency as compared to the mutations on *haplotigs* as the reads of the non-mutated alleles were aligned in this region as well) (Fig. 1f).

To understand the layer-specific dynamics of gene conversions, we also analyzed the allele frequencies of the converted alleles within the samples where the conversions had not been observed. It is important to note that our approach could not detect gene conversions that removed alleles from specific layers while the converted alleles still remained in other layers. Here we could only find those gene conversions that removed alleles entirely (either from all or from few layers).

We found one predominant cluster where the allele frequencies of the converted alleles were intermediate in the *L_12_* samples and low (but not absent) in the *L_123_* samples (Fig. 1f; Supplementary Table S2). This suggested that the heterozygous alleles were present only in *L_1_* or *L_2_* before they got converted. As for DNM, highly similar patterns in allele frequencies regarding gene conversions were found on *haplotigs* and on *homotigs* (Fig. 1f).

In summary, approximately one third of the *de novo* mutations occurred in specific meristematic layers only, while two third were found in cells derived from all layers. In contrast we did not find any gene conversion that would remove the allele from all cell-layers (i.e., a case with 50% alternative allele frequency in all wildtype samples while 0% alternative allele frequency in all mutant samples) implying that all of them were layer-specific.

### Distribution of somatic mutations across individual branches

The samples we sequenced to identify somatic mutations were based on bulked leaves sampled from scaffolds of the upper (mutant) and lower (wildtype) part of the tree (Fig. 1a). To get a better understanding of the distribution of the mutations across the tree, we checked the mutant and wildtypes alleles of the 29 mutations within the Sanger-sequencing data that we used for evaluating the mutation identification workflow (including 20 gain-of-heterozygosity and 9 loss-of-heterozygosity separating the mutant and wildtype parts). The individual leaves used for the genotyping were sampled from seven upper and six lower scaffolds of the tree (Fig. 1a; Material and Methods). Besides individual leaf samples, we also tested the existence of these mutations in pooled DNA, including two leaf samples and two petiole samples for the mutant part, and similarly for the wildtype part. In total, each mutation was genotyped in 21 DNA samples.

Among the 20 gain-of-heterozygosity mutations (Supplementary Figure S2-21), we found 17 in all seven mutant scaffolds of the tree and not in any of the lower wildtype scaffold samples (Supplementary Figure S2-4,6-7,9-20). Among the remaining three, one was found in six mutant scaffolds while the Sanger sequencing related to the other mutant scaffold was not successful (Supplementary Figure S5); one was also found in six mutant scaffolds plus one lower wildtype scaffold (Supplementary Figure S8); while the last one could only be found in pooled samples of the upper mutant scaffolds (Supplementary Figure S21). The nine loss-of-heterozygosity mutations were all found in two to six samples of the lower wildtype part of the tree but not in any of the samples of the upper part of the tree (Supplementary Figure S22-30).

### Identification of candidate genes causing enhanced red fruit skin coloration

To understand the molecular basis of the coloration differences between the upper mutant and the lower wildtype parts of the *RubyMac* tree, using the RNA-seq data as mentioned above, we first analyzed genome-wide expression differences as they occur in the skins of mature mutant and wildtype apples. This revealed 56 differentially expressed genes (DEG), including 48 of them being up-regulated in the mutant fruits (Fig. 2a; Materials and Methods). Apart from putative disease resistance and uncharacterized genes, 18 of the DEGs (following the GDDH reference genome annotation) were involved in flavonoid or anthocyanin biosynthesis and regulation, including the *MdMYB*1 transcription factor *MD09G1278600* that regulates apple skin color^20–24^ (Supplementary Table S3; Materials and Methods). While this already indicates the alteration of the pathways involved in the production of fruit coloration, we could not identify any genomic mutations within the DEGs or in their flanking regions (up to hundreds of kb) suggesting that the expression differences are downstream effects of a mutation somewhere else in the genome^55^.

We examined the 53 somatic changes that discriminated the mutant and the wildtype parts for their effects on gene integrity (consisting of 22 DNM, of which all occurred in all three layers, and 31 gene conversions). Of those, 12 mutations were located in genes, including four nonsynonymous and one synonymous change in CDS, five in introns, and two in UTRs (Fig. 2b; Supplementary Table S2). This revealed several candidate genes (Fig. 1c; Supplementary Table S4-6), including a non-synonymous mutation in an F-box gene (at Chr09:32,528,560, when mapped to the GDDH13 v1.1 genome assembly) which was within a GWAS peak that was recently associated with red skin coloration^5–6,37–38^ as well as a mutation at Chr09:4,621,163, which was 43 bp away from an anthocyanins 5-aromatic acyltransferase-like protein and near a QTL that was identified as involved red skin coloration variation^37^. Another nonsynonymous mutation was found in a homolog of the *Arabidopsis thaliana HSP90* gene (MD08G1011200), which has been shown to influence morphogenetic responses to environmental stresses such as leaf colors^75^.

## DISCUSSION

Somatic mutations can lead to immediate phenotypic consequences. Specifically, mutations that occur in the meristem have the chance to be propagated into large sectors and thereby drastically change the behavior of large parts of a plant. In fruit trees, somatic mutations were the basis for new traits and various of those have been introduced into different breeding programs. Knowing the genetic basis of bud sport mutations can broaden our understanding of phenotypic change and accelerate breeding related efforts. Analyzing the different sectors of an apple tree with a sport mutation that changed fruit coloration, we identified somatic mutations that distinguished the wildtype and mutant genomes and thereby we were able to define candidate mutations that might underly the change in fruit coloration.

Unexpectedly, however, we did not only find gain-of-heterozygosity mutations, but also a similar number of loss-of-heterozygosity mutations. To investigate the general differences between them, we investigated the meristematic layers using tissue-specific DNA samples that were enriched for cells derived from specific layers. *De novo* mutations were observed either to result from *L_3_* or from all layers. Surprisingly, however, we could hardly find any *de novo* mutations that were specific to *L_1_* or *L_2_*. While *L_1_*-specific mutations might have been lost due to an underrepresentation of *L_1_* cells in the sequenced samples, *L_2_* mutations were apparently occurring at a much lower frequency as compared to *L_3_* specific mutations. This is remarkable, as meiotic precursor cells (as well as all gamete cells) result from *L_2_* and therefore specifically the mutations in *L_2_* have the chance to be passed on the next generation. In contrast, loss-of-heterozygosity mutations were specific to *L_1_* or *L_2_* or both.

Moreover, somatic *de novo* mutations were enriched for GC=>AT transitions, which is consistent with the spectrum of germline mutations in plants^25^, while loss-of-heterozygosity mutations were biased towards AT=>GC transitions. This suggested that the genome-wide GC-content might be controlled by both novel mutations and the mismatch repairing systems, which can counteract each other’s biases. It should be noted, we cannot fully exclude the possibility that the loss-of-heterozygosity mutations were in fact *de novo* mutations that occurred only in the lower part of the tree. But if we assume that the gene conversions were wrongly assigned (and were in fact *de novo* mutations), we would expect them in similar layers as all other *de novo* mutations, which, however, was not the case (gene conversions were specific to *L_1_* or *L_2_* while most of the layers-specific mutations were found in *L_3_*). Another note of caution, in our interpretation we have assumed a three-layer model of the meristem, as well as the persistency of the layer identify in somatic tissue. While both might be a simplification with respect to the apple tree analyzed here, our analysis does confirm that layer-specificity in general needs to be considered when analyzing somatic mutations in plants and that the search for mutations that are fixed in one haplotype would have missed some of the somatic mutations.

Future application of single cell sequencing technologies might help to overcome some of the challenges presented and help to identify differences in the mutational profiles of different layers and thereby help deepen our understanding of the dynamics of somatic mutations and their inheritance in long-lived plants.

## Materials and Methods

### DNA/RNA extraction and sequencing library preparation

Trichome tissue (*L_1_*) was separated by freezing mutant and wildtype leaves in liquid nitrogen and dislodging the trichomes into a 50-mL Falcon tube using a soft paintbrush. The liquid nitrogen was allowed to boil off leaving the trichomes behind. Leaf surface tissue (*L_12_*) was isolated from mutant and wildtype leaf petioles by scraping the petiole with a razor blade to remove the outer tissue layers and directly drop them into liquid nitrogen and stored at −80 °C until use. Whole leaves (*L_123_*) as well *L_12_* tissue was ground in liquid nitrogen in a mortar and pestle prior to extraction. DNA was extracted using the DNeasy Plant Mini Kit (Qiagen, Hilden, Germany). DNA samples of whole leaves (*L_123_*) were used for both Illumina and Sanger sequencing.

Fruit skin from the mutant fruit and wildtype fruit (each with three replicates) was respectively obtained using a handheld fruit peeler to separate the skin (as well as a few millimeters of cortex tissue) from the fruit. Tissue was ground in liquid nitrogen using a mortar and pestle and stored at −80 °C until use. Total RNA was extracted from according to Gasic et al. (2004)^61^ and adapted for microcentrifuge tubes by using 1/10 the recommended volumes of all solvents and sample weights. Following the initial extraction, the extracted RNA was purified using the RNeasy mini-spin column protocol (Qiagen, Hilden, Germany).

All sequencing libraries were prepared at Max Planck-Genome-center Cologne, Cologne, Germany. Twelve DNA libraries were sequenced using Illumina HiSeq2500/MiSeq platforms under 250~300 bp paired-end reads mode. Six RNA libraries (mutant with three replicates and wildtype with three replicates) were sequenced using Illumina HiSeq3000 under 150 bp single-end reads mode.

### Genome size estimation

For each of the eight *L_12_* and *L_123_* samples, *k*-mer counting (*k*=21) in the respective short reads was performed using the tool *jellyfish*^62^. Given the *k*-mer frequency histograms, genome size and rate of heterozygosity were estimated using *findGSE*^41^ under heterozygous mode. Averages of the eight estimations (with standard variations) were defined as the final value.

### Genome assembly and annotation

Genome assembly was performed using *DiscovarDeNovo*^34^ with default settings, which led to the *RubyMac DDN* genome assembly of 875 Mb including all ≥1 kb contigs. For gene prediction, Illumina single-end 150 bp RNA reads were aligned to the assembly using *tophat2*^44^ with options -N (read mismatches) and --read-edit-dist as 10, --library-type fr-firststrand and the others as defaults. The result BAM file was provided to *AUGUSTUS*^45^ *bam2hints* to generate a GFF file for guiding gene annotation by *AUGUSTUS* with options --species=Arabidopsis --extrinsicCfgFile=/augustus-master/config/extrinsic/extrinsic.cfg --alternative-from-evidence=true --UTR=on --progress=true -- allow_hinted_slicesites=atac --uniqueGeneId=true.

### Haplotig identification

To guarantee a sufficient sequencing depth required by this analysis, paired-end reads from two wildtype *L_123_* samples were merged as one set, each contributing ~75x (Supplementary Table S1). Combined reads were aligned to the *RubyMac DDN* assembly using *minimap2*^63^, and the bam file was sorted and indexed using *samtools*^64^. Then “*purge_haplotigs readhist*” was applied to generate a sequencing depth histogram for each contig. Expecting ~150x sequencing depth for a homotig and ~75x for a haplotig, we set 20, 122, and 300 as depth cutoffs required by *Purgen_Haplotigs*, corresponding to low (“valley” on the left of the heterozygous peak in the depth histogram), middle (“valley” between the heterozygous and homozygous peaks) and high (nearly two times of the homotig sequencing depth). With these cutoffs, “*purge_haplotigs contigcov*” would predict a contig as a haplotig if more than 30% of its positions were not covered by 122~300x, for which “*purge_haplotigs purge*” would search for a matching haplotig and if 85% of the contig can be aligned with others, it would be determined as haplotig. This analysis led to 59,293 curated primary contigs (~630.7 Mb), 44,305 haplotigs (~242.2 Mb) and 795 artificial contigs (~1.8 Mb).

### Small-scale variation identification and validation with Sanger sequencing

We first identified small-scale variations with a customized pipeline using the *RubyMac DDN* assembly as reference. Specifically, reads of each of the 12 DNA samples were respectively aligned to the reference genome using *bowtie2*^65^, and resulted SAMs was converted to BAMs and indexed using *samtools*^64^. With each BAM, read counts were collected for all bases (including deletions) at each position of the reference genome by “*shore consensus*”^66^, which simultaneously predicted variations with allele frequency, read coverage and base quality score (maximum 40). Then a somatic mutation would be called at a position, if there was an alternative allele not found in all eight *L_12_* and *L_123_* samples (note that due to the limited sequencing depth of *L_1_* samples, they were not considered in this step).

With this, we selected the initial list of 69 mutations for Sanger sequencing validation, where the alternative allele must be covered by at least 15 reads in at least one sample and not common between wildtype and mutant branches of the tree. The initial list of 69 mutant- or wildtype-specific mutations were genotyped in DNA of individual leaves from seven mutant scaffolds on the tree, two pooled leaf blade samples (mutant scaffolds {5, 7} and {4, 6}) and two pooled petiole samples (mutant scaffolds {5, 7} and {4, 6}), and similarly in DNA of individual leaves from six wildtype scaffolds on the tree, two pooled leaf blade samples (wildtype scaffolds {5, 7} and {4, 6}) and two pooled petiole samples (wildtype scaffolds {5, 7} and {4, 6}), with 19-29 bp primers around the mutations designed by *Primer3*^68^ (Fig. 1a; Supplementary Table S2). Generally, each mutation was genotyped in 21 DNA samples. The R package ‘*sangerseqR*’ was used to analyze the Sanger-seq data with the threshold of 0.01 for function *makeBaseCalls* to call (low-frequency) alternative alleles^77^.

In the second round of mutation identification, the following criteria were applied in each somatic mutation calling, including minimum alternative allele frequency of 0.1, read coverage of 10~150x, minimum alternative base quality of 30 (or Q300. Note, if there were >2 samples with the alternative allele, the mutation was kept only if at least two samples showing Q30. Otherwise, the mutation was kept if at least one sample showing Q30, and both the maximum average numbers of unknown base *N* and mismatches in the read alignments covering the mutation position were set as 5 (corresponding to ~2% of the read length of 250~300 bp). The detected variations were further filtered manually by checking alignments by browsing with IGV^67^. This led to the list of 100 mutations.

In addition, the reported variations (Supplementary Table S2) could also be detected by other tools such as reference-based *MuTect* (version 1.1.4)^36^ with options *“--input_file:normal sample1.bam --input_file:tumor sample2.bam*”, where sample1.bam and sample2.bam are BAM files as mentioned above, and reference-free *discosnp*++ (version 2.2.X)^46^ with options “*-r readset.txt -k 41 -b 1*”, where the file readset.txt included the list of fastq files of any pair of samples to compare.

### Somatic mutation clustering

Let *r_123_* and *a_123_* respectively represent for the counts of reads carrying reference and alternative alleles for *L_123_* (i.e., blade sample) while *r_12_* and *a_12_* for *L_12_* (i.e., surface sample) in either mutant or wildtype, depending on which one has the alternative allele. The difference between (*r_123_*, *a_123_*) and (*r_12_*, *a_12_*) was examined by a two-sided *Fisher*’s exact test (*R* package “stats”). To reduce the effect of randomness in read counts, any value in the set of (*r_123_*, *a_123_*, *r_12_*, *a_12_*) for one mutation was taken as the summation of read counts from two replicates. If the test for a mutation gives a *p*- value smaller than 0.05, it was defined as with significant difference in allele frequencies (AFs) of *L_123_* and *L_12_*, indicating there was a gain (or loss) of the alternative allele in a layer. This led to three clusters of mutations (on haplotigs and homotigs), a) AF in *L_123_* is comparable to *L_12_*, b) AF in *L_123_* is significantly larger than *L_12_*, and c) AF in *L_12_* is significantly larger than *L_123_*.

### Differential gene expression analysis

The RNA-seq reads of three mutant-related replicates (upper part of the tree) and three wildtype-related replicates (lower part of the tree) were respectively aligned to the 875 Mb *RubyMac DDN* genome using *tophat2*^44^, with options “-p 10 -a 10 -g 10 --library-type fr-firststrand” for controlling the number of threads, minimum anchored bases and maximum multiple hits of reads. The BAM files were indexed with *samtools*^64^. Read counts for genes in GFF (generated by *AUGUSTUS*^45^) were extracted with *HTSeq*^69^ for each sample. Differential expression (DE) analysis was performed with an *R* pipeline^70^, with adjusted *p*-value<0.05 for considering differential expression.

We also repeated the analysis with the 807 Mb GDDH13 v1.1 reference genome and annotation of *Golden Delicious*^40^. Four DE genes as identified with the *RubyMac DDN* assembly (for which blasts to the NCBI nucleotide databases hit genes of probable copper-transporting ATPase *HMA5*, not available, F-box protein *SKIP27*-like and myosin heavy chain clone 203-like) were not in *Golden Delicious* gene annotation. To be comprehensive, we created a final list of 56 DE genes by merging two sets (Supplementary Table S3).

### Large indels and copy number variation (CNV) identification

The ~875 Mb *RubyMac DDN* assembly was indexed with *bowite2*^65^. Reads were aligned for each of the eight *L_12_* and *L_123_* samples with *bowite2*^65^, and the BAM files were indexed with *samtools*^64^. Then the BAM files were provided to *CNVnator* (version)^60^ for CNV detection with various bin sizes of 50, 200 and 300 bp. After filtering, we could find 199 mutant-specific and 169 wildtype-specific candidate CNVs. However, further checking of the candidates on alignments in IGV did not reveal any convincing ones.

## Data availability

Read data of all 12 DNA sequencing libraries, 6 RNA sequencing libraries, and Sanger sequencings of 69 mutations within 21 samples are available under BioProject PRJNA881844 at NCBI.

## Acknowledgements

The authors would like to thank Saurabh Pophaly (MPIPZ) for help in data management. Randy Beaudry acknowledges support from Michigan AgBioResearch and the USDA National Institute of Food and Agriculture, Hatch project MICL002688, and financial support from the Michigan Apple Committee. Maria von Korff and Korbinian Schneeberger acknowledge the Deutsche Forschungsgemeinschaft (DFG, German Research Foundation) under Germany′s Excellence Strategy—EXC-2048/1—Project ID: 390686111.

## Author contributions

K.S. and R.B. designed and supervised the project. H.S., J.C., W.J., E.W. and K.S. performed all sequence analysis. P.A., R.B, T.R. M.K. and B.H. performed all wet-lab experiments. H.S., R.B. and K.S. wrote the manuscript. All authors read and approved the final manuscript.

