## Supplementary Figures for "The identification and analysis of meristematic mutations within the apple tree that developed the *RubyMac* sport mutation"

Sun *et al. 2022*

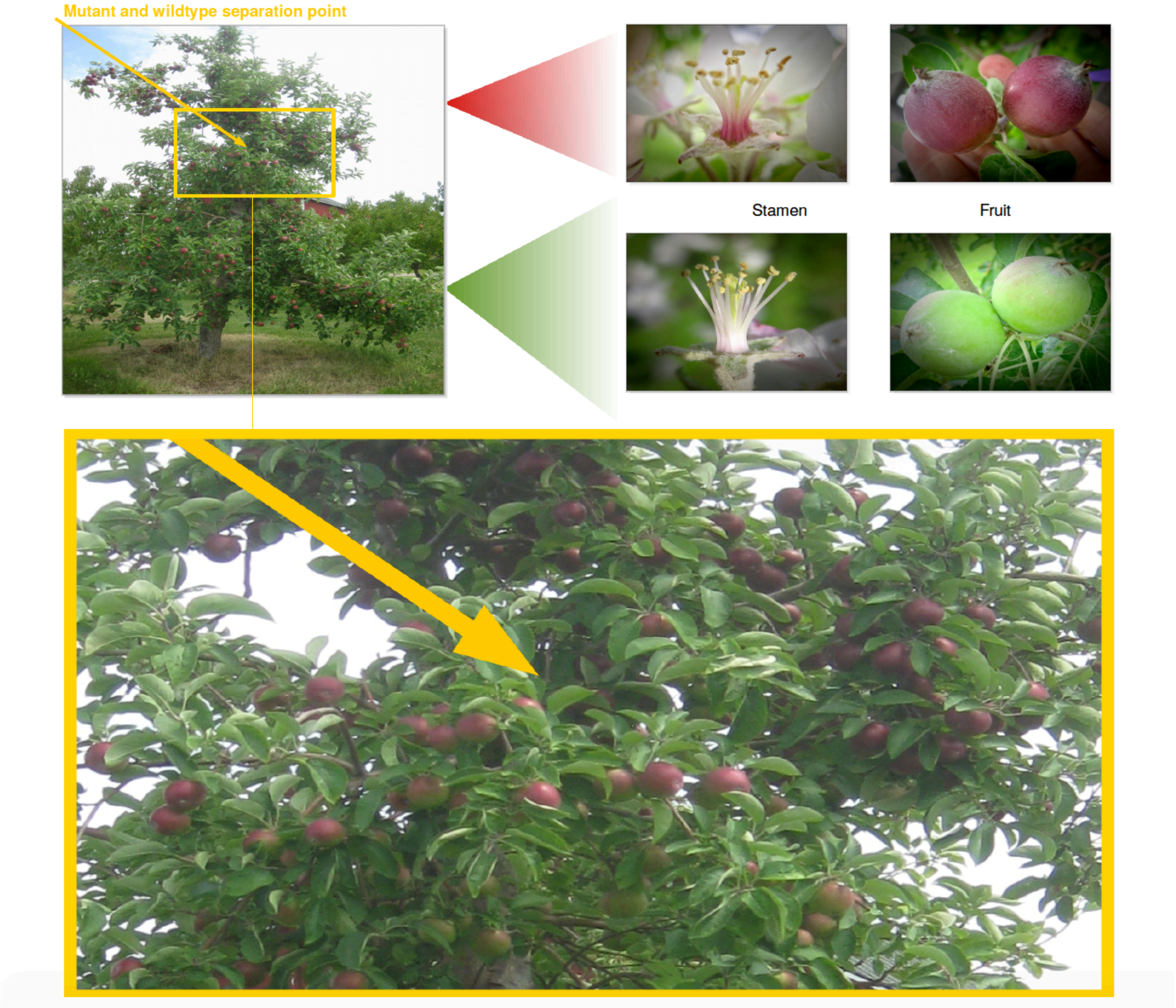


**Supplementary Figure 1. Phenotype of the sport *RubyMac* tree.**

The tree is growing at Michigan, USA. Phenotypically, the seven younger (higher) and ten older (lower) scaffolds, as separated by the site pointed by the arrow in yellow, were identified as the mutant and the wildtype. The rectangle in yellow shows a zoom into the tree section containing the separation point. As shown, both the stamen at an early stage and fruits at a late stage of fruit development showed clearly higher levels of red/purple pigment accumulation than the wildtype branches.

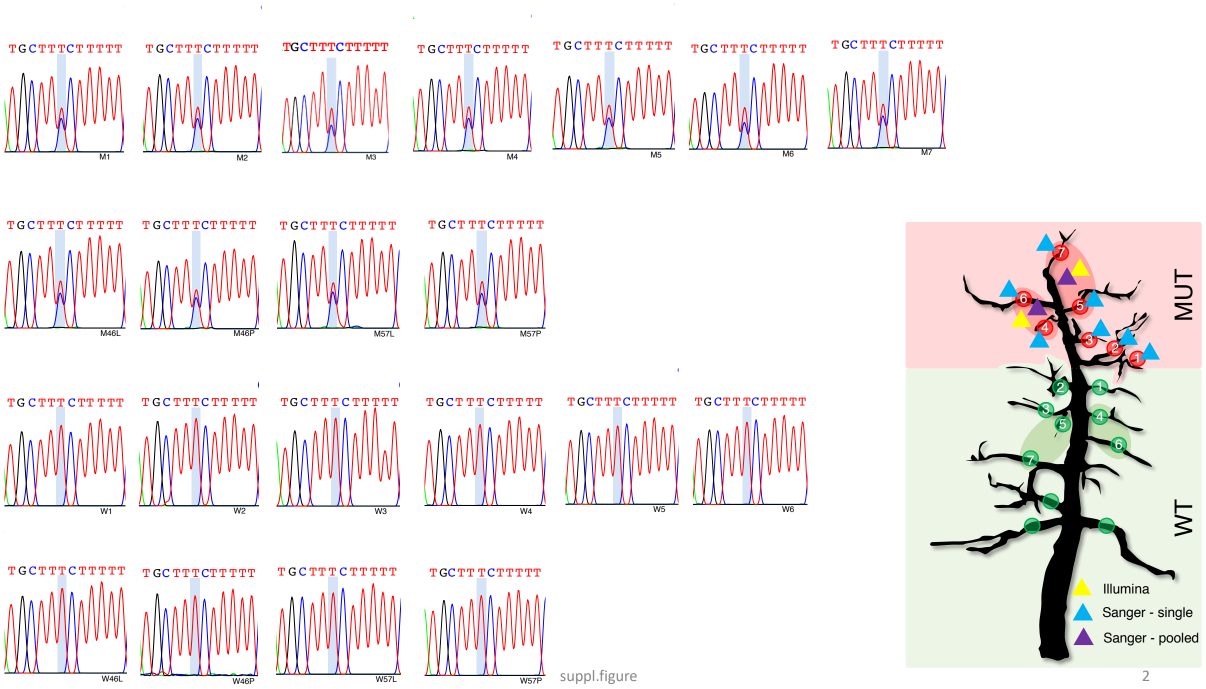


**Supplementary Figure 2. Sanger-seq of mutation No. 1 on fl_10005 (Supplementary Table S2).** MUT: mutant, WT: wildtype. The mutation was found at mutant scaffolds MUT1-7 (as indicated by triangles in blue in the tree sketch), in pooled petiole MUT46P/57P and leaf MUT46L/57L (triangles in purple), while it was not found in any of the individual/pooled wildtype scaffolds. Triangles in yellow indicated the mutations was found by Illumina sequencing of pooled leaf DNA of {4,6} or {5, 7} scaffolds. The mutation site was indicated by the vertical bar in light blue in each Sanger sequencing sample.

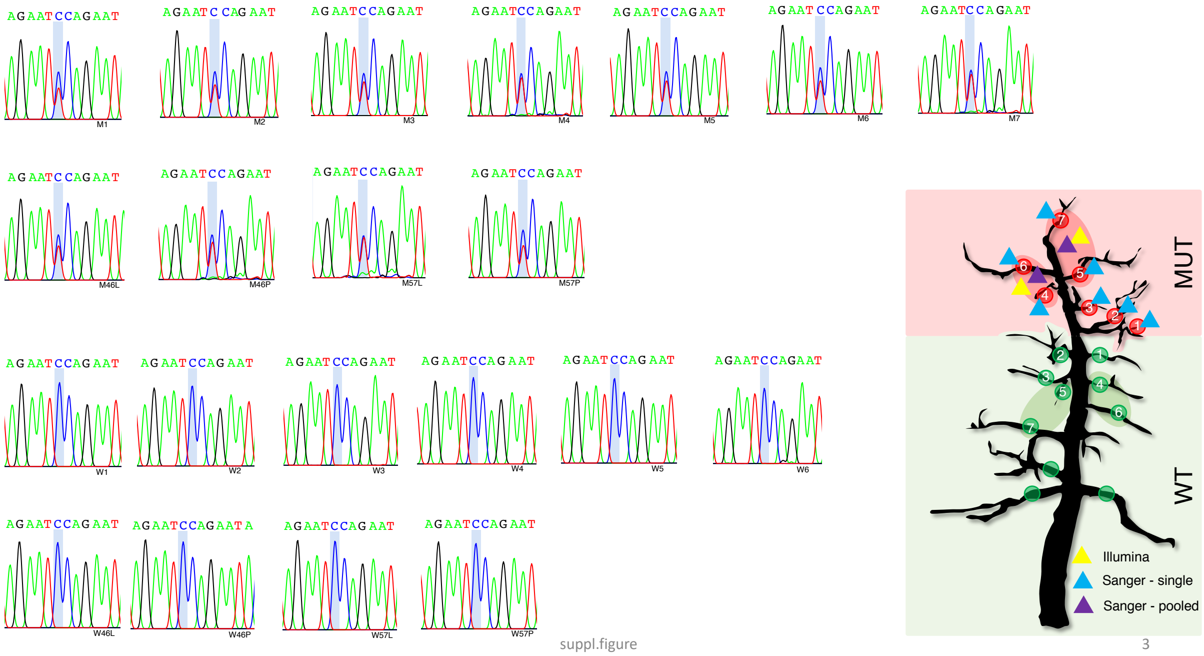


**Supplementary Figure 3. Sanger-seq of mutation No. 2 on fl_10473 (Supplementary Table S2).** MUT: mutant, WT: wildtype. The mutation was found at mutant scaffolds MUT1-7 (as indicated by triangles in blue in the tree sketch), in pooled petiole MUT46P/57P and leaf MUT46L/57L (triangles in purple), while it was not found in any of the individual/pooled wildtype scaffolds. Triangles in yellow indicated the mutations was found by Illumina sequencing of pooled leaf DNA of {4,6} or {5, 7} scaffolds. The mutation site was indicated by the vertical bar in light blue in each Sanger sequencing sample.

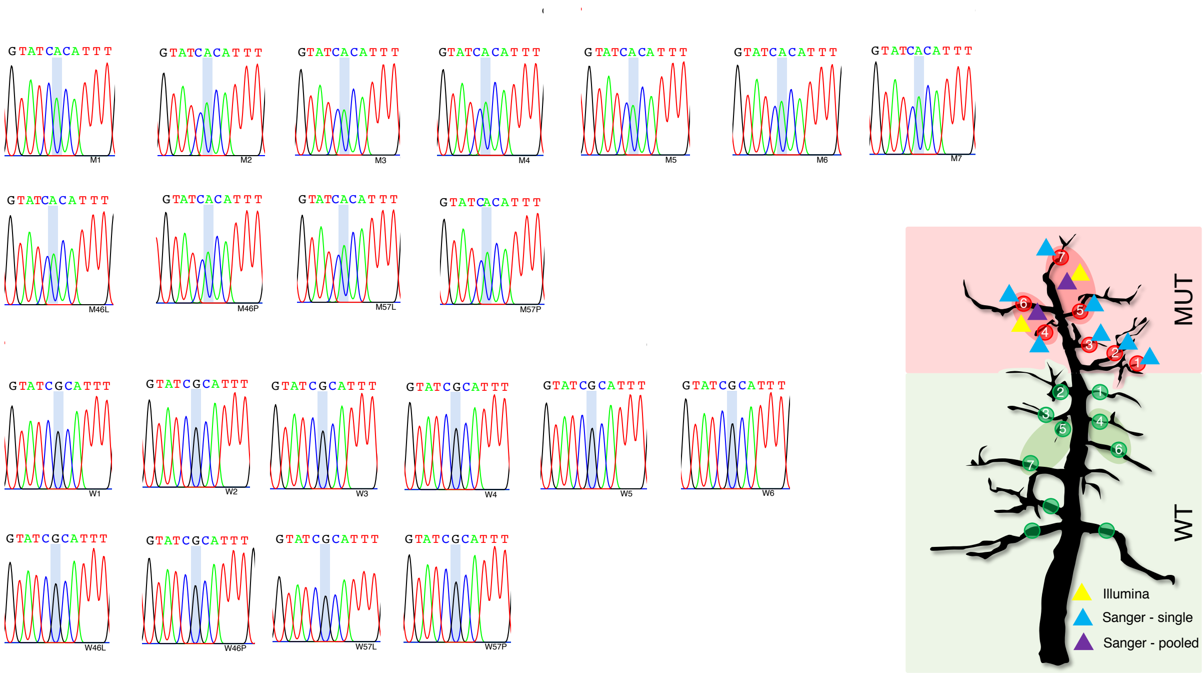


suppl.figure 4

**Supplementary Figure 4. Sanger-seq of mutation No. 5 on fl_11153 (Supplementary Table S2).** MUT: mutant, WT: wildtype. The mutation was found at mutant scaffolds MUT1-7 (as indicated by triangles in blue in the tree sketch), in pooled petiole MUT46P/57P and leaf MUT46L/57L (triangles in purple), while it was not found in any of the individual/pooled wildtype scaffolds. Triangles in yellow indicated the mutations was found by Illumina sequencing of pooled leaf DNA of {4,6} or {5, 7} scaffolds. The mutation site was indicated by the vertical bar in light blue in each Sanger sequencing sample.

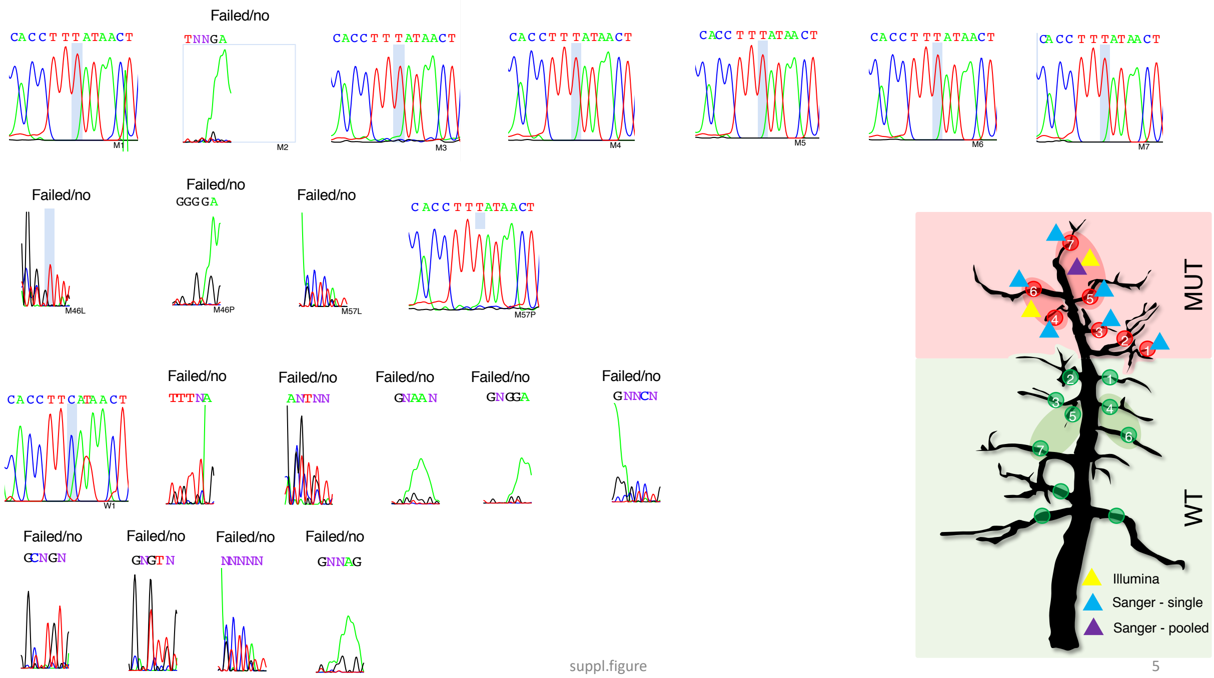


**Supplementary Figure 5. Sanger-seq of mutation No. 9 on fl_12199:C$\rightarrow$T (Supplementary Table S2).** MUT: mutant, WT: wildtype. The mutation was found at mutant scaffolds MUT1,3-7 (as indicated by triangles in blue in the tree sketch), in pooled petiole MUT57P (triangles in purple), while it was not found in the individual wildtype scaffold WT1. Triangles in yellow indicated the mutations was found by Illumina sequencing of pooled leaf DNA of {4,6} or {5, 7} scaffolds. The mutation site was indicated by the vertical bar in light blue in each Sanger sequencing sample. "Failed/no" indicated cases where Sanger sequencing did not work or no alternative allele could be found.

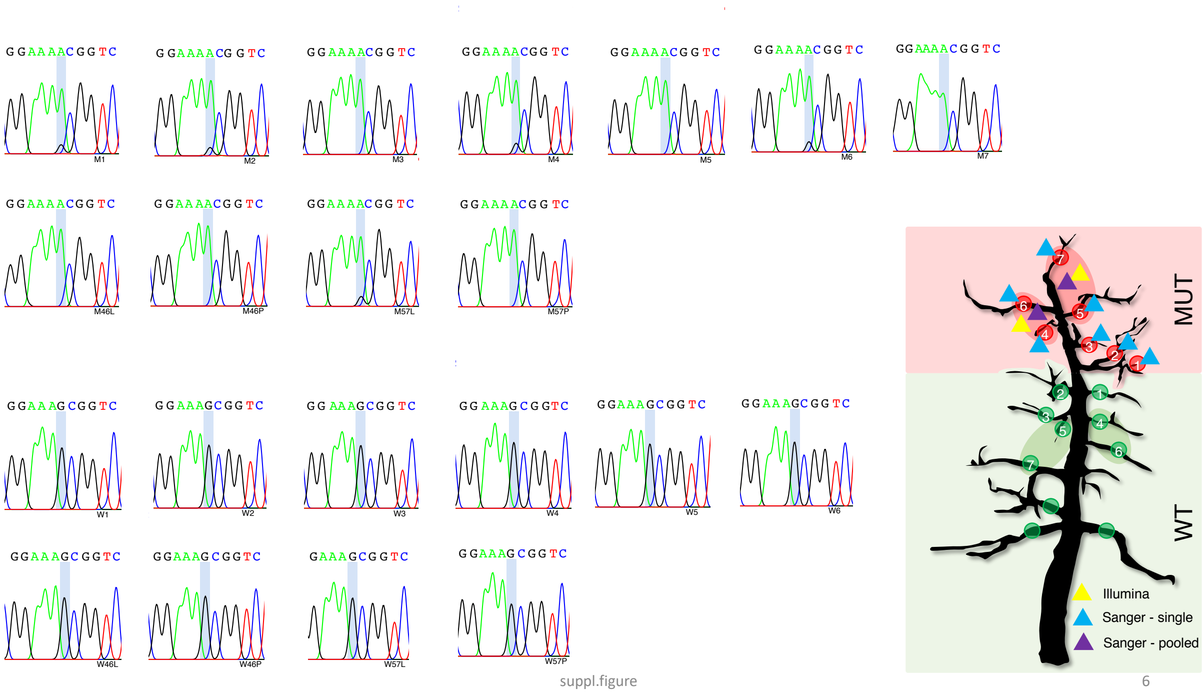


**Supplementary Figure 6. Sanger-seq of mutation No. 12 on fl_12691: G→A (Supplementary Table S2).** MUT: mutant, WT: wildtype. The mutation was found at mutant scaffolds MUT1-7 (as indicated by triangles in blue in the tree sketch), in pooled petiole MUT46P/57P and leaf MUT46L/57L (triangles in purple), while it was not found in any of the individual/pooled wildtype scaffolds. Triangles in yellow indicated the mutations was found by Illumina sequencing of pooled leaf DNA of {4,6} or {5, 7} scaffolds. The mutation site was indicated by the vertical bar in light blue in each Sanger sequencing sample.

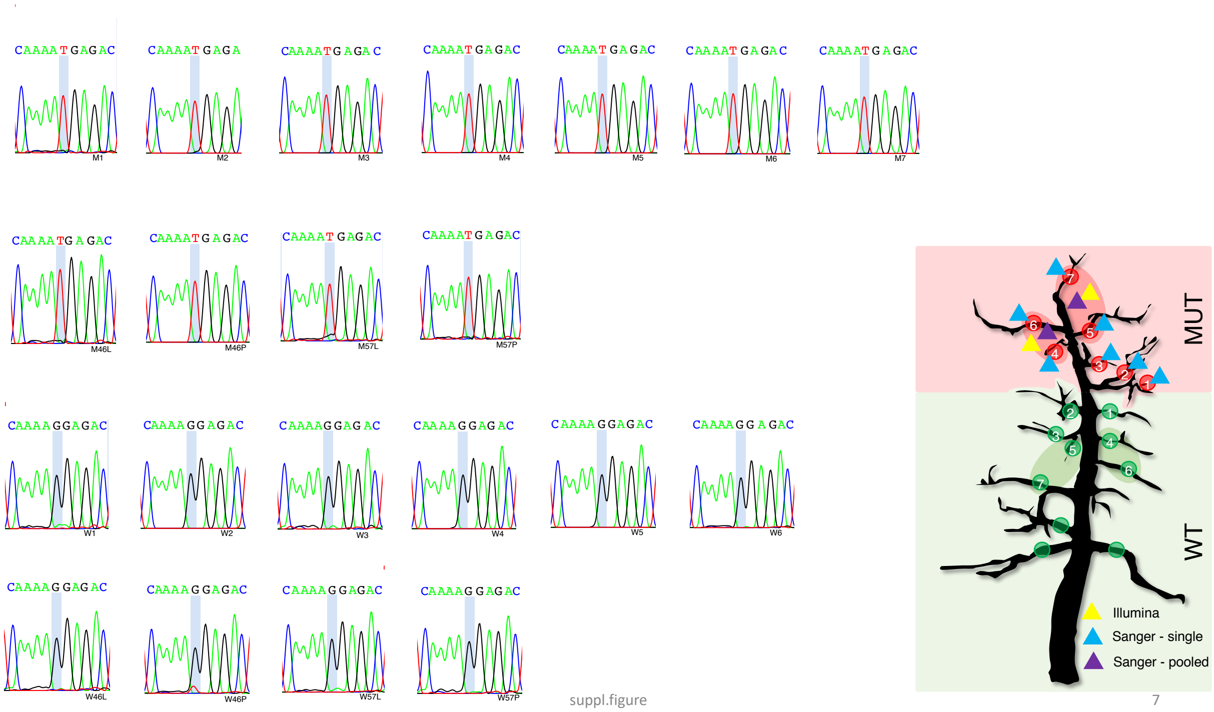


suppl.figure 7

**Supplementary Figure 7. Sanger-seq of mutation No. 23 on fl_1690:G→T (Supplementary Table S2).** MUT: mutant, WT: wildtype. The mutation was found at mutant scaffolds MUT1-7 (as indicated by triangles in blue in the tree sketch), in pooled petiole MUT46P/57P and leaf MUT46L/57L (triangles in purple), while it was not found in any of the individual/pooled wildtype scaffolds. Triangles in yellow indicated the mutations was found by Illumina sequencing of pooled leaf DNA of {4,6} or {5, 7} scaffolds. The mutation site was indicated by the vertical bar in light blue in each Sanger sequencing sample.

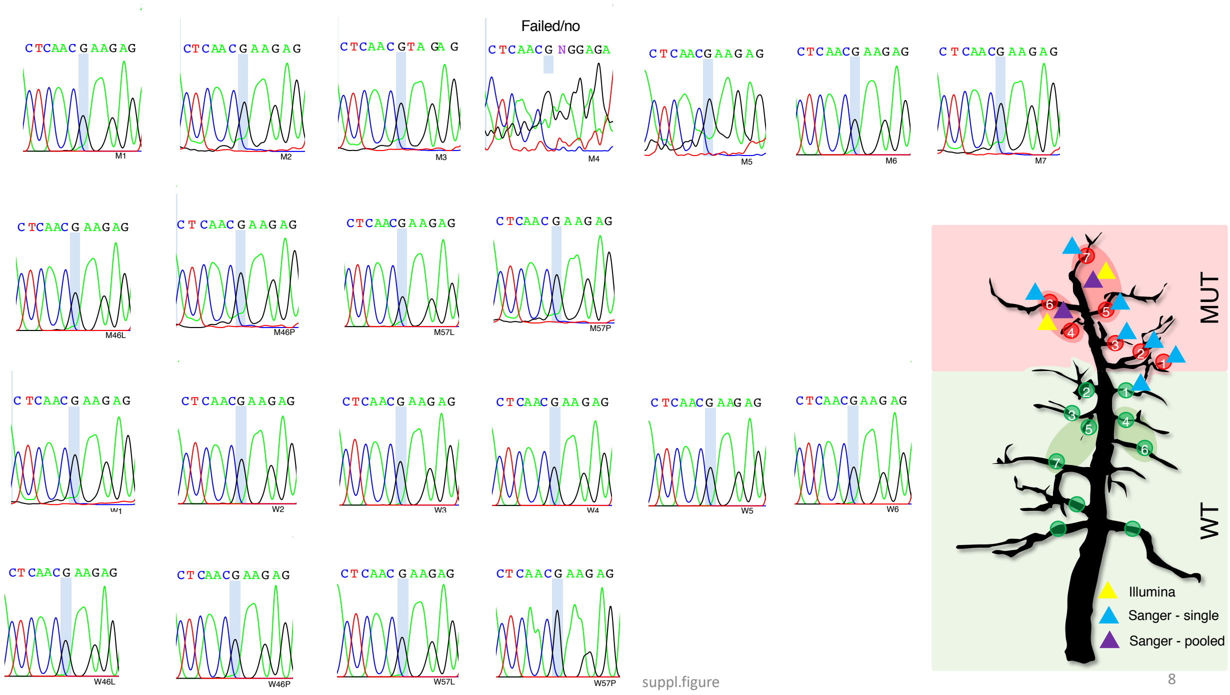


**Supplementary Figure 8. Sanger-seq of mutation No. 24 on fl_16917:G→A (Supplementary Table S2).** MUT: mutant, WT: wildtype. The mutation was found at mutant scaffolds MUT1-3, 5-7 (as indicated by triangles in blue in the tree sketch), in pooled petiole MUT46P/57P and leaf MUT46L/57L (triangles in purple), while, except for WT1, it was not found in any of the other individual/pooled wildtype scaffolds. Triangles in yellow indicated the mutations was found by Illumina sequencing of pooled leaf DNA of {4,6} or {5, 7} scaffolds. The mutation site was indicated by the vertical bar in light blue in each Sanger sequencing sample. “Failed/no” indicated cases where Sanger sequencing did not work or no alternative allele could be found.

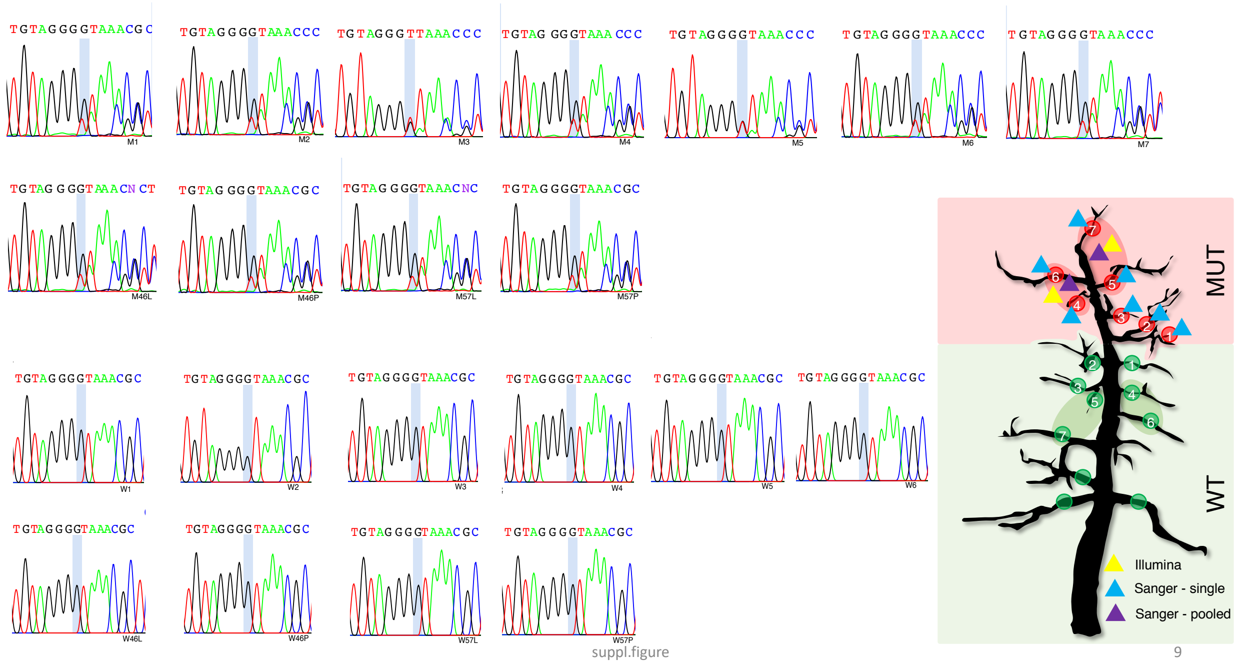


**Supplementary Figure 9. Sanger-seq of mutation No. 25 on fl_169395:G→Deletion (Supplementary Table S2).** MUT: mutant, WT: wildtype. The mutation was found at mutant scaffolds MUT1-7 (as indicated by triangles in blue in the tree sketch), in pooled petiole MUT46P/57P and leaf MUT46L/57L (triangles in purple), while it was not found in any of the individual/pooled wildtype scaffolds. Triangles in yellow indicated the mutations was found by Illumina sequencing of pooled leaf DNA of {4,6} or {5, 7} scaffolds. The mutation site was indicated by the vertical bar in light blue in each Sanger sequencing sample.

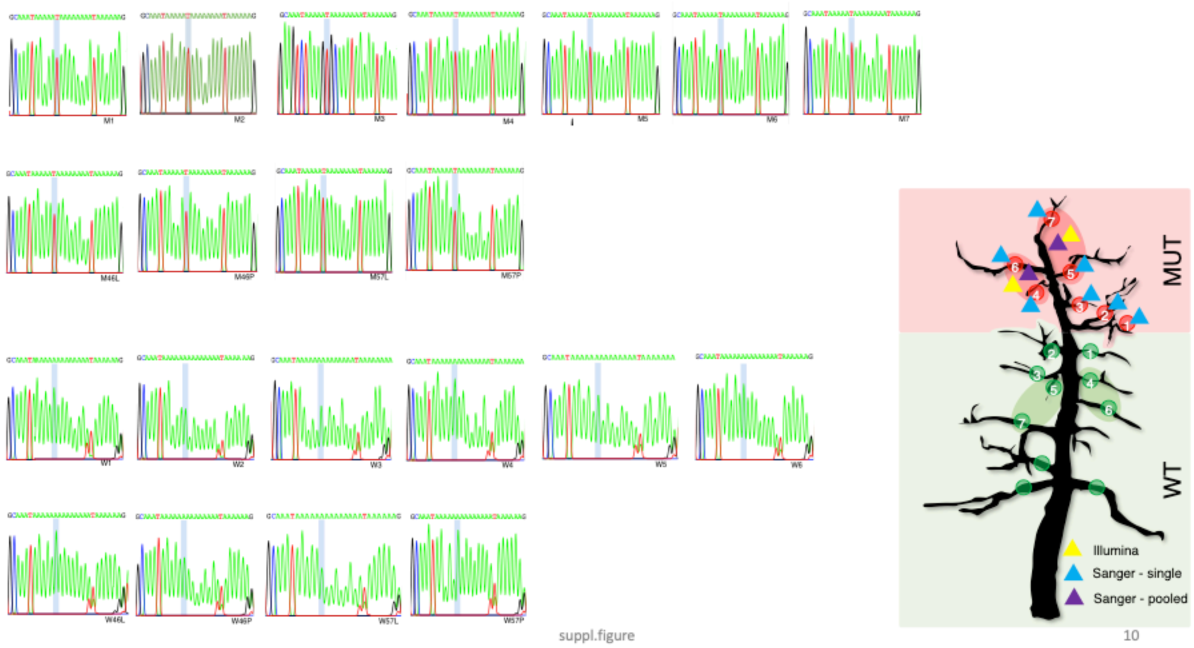


**Supplementary Figure 10. Sanger-seq of mutation No. 30 on fl_20119:A→T (Supplementary Table S2).** MUT: mutant, WT: wildtype. The mutation was found at mutant scaffolds MUT1-7 (as indicated by triangles in blue in the tree sketch), in pooled petiole MUT46P/57P and leaf MUT46L/57L (triangles in purple), while it was not found in any of the individual/pooled wildtype scaffolds. Triangles in yellow indicated the mutations was found by Illumina sequencing of pooled leaf DNA of {4,6} or {5, 7} scaffolds. The mutation site was indicated by the vertical bar in light blue in each Sanger sequencing sample.

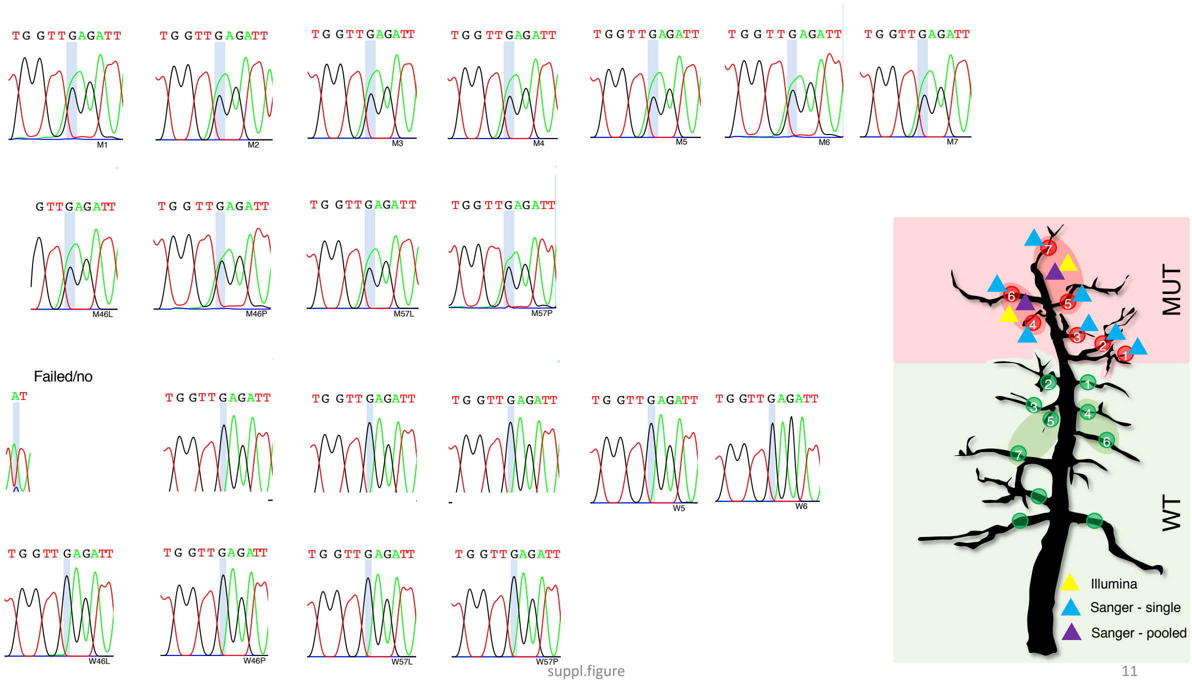


**Supplementary Figure 11. Sanger-seq of mutation No. 33 on fl_218:G→A (Supplementary Table S2).** MUT: mutant, WT: wildtype. The mutation was found at mutant scaffolds MUT1-7 (as indicated by triangles in blue in the tree sketch), in pooled petiole MUT46P/57P and leaf MUT46L/57L (triangles in purple), while it was not found in any of the individual/pooled wildtype scaffolds. Triangles in yellow indicated the mutations was found by Illumina sequencing of pooled leaf DNA of {4,6} or {5, 7} scaffolds. The mutation site was indicated by the vertical bar in light blue in each Sanger sequencing sample. “Failed/no” indicated cases where Sanger sequencing did not work or no alternative allele could be found.

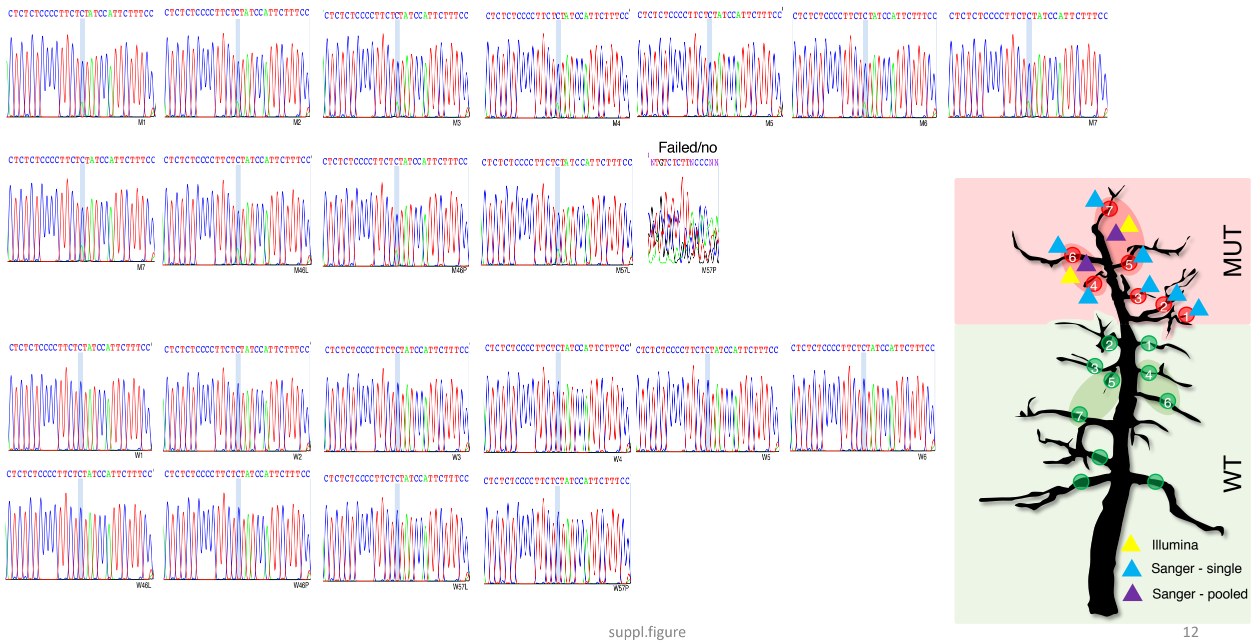


**Supplementary Figure 12. Sanger-seq of mutation No. 44 on fl_29360: C→A (Supplementary Table S2).** MUT: mutant, WT: wildtype. The mutation was found at mutant scaffolds MUT1-7 (as indicated by triangles in blue in the tree sketch), in pooled petiole MUT46P and leaf MUT46L/57L (triangles in purple), while it was not found in any of the individual/pooled wildtype scaffolds. Triangles in yellow indicated the mutations was found by Illumina sequencing of pooled leaf DNA of {4,6} or {5, 7} scaffolds. The mutation site was indicated by the vertical bar in light blue in each Sanger sequencing sample. “Failed/no” indicated cases where Sanger sequencing did not work or no alternative allele could be found.

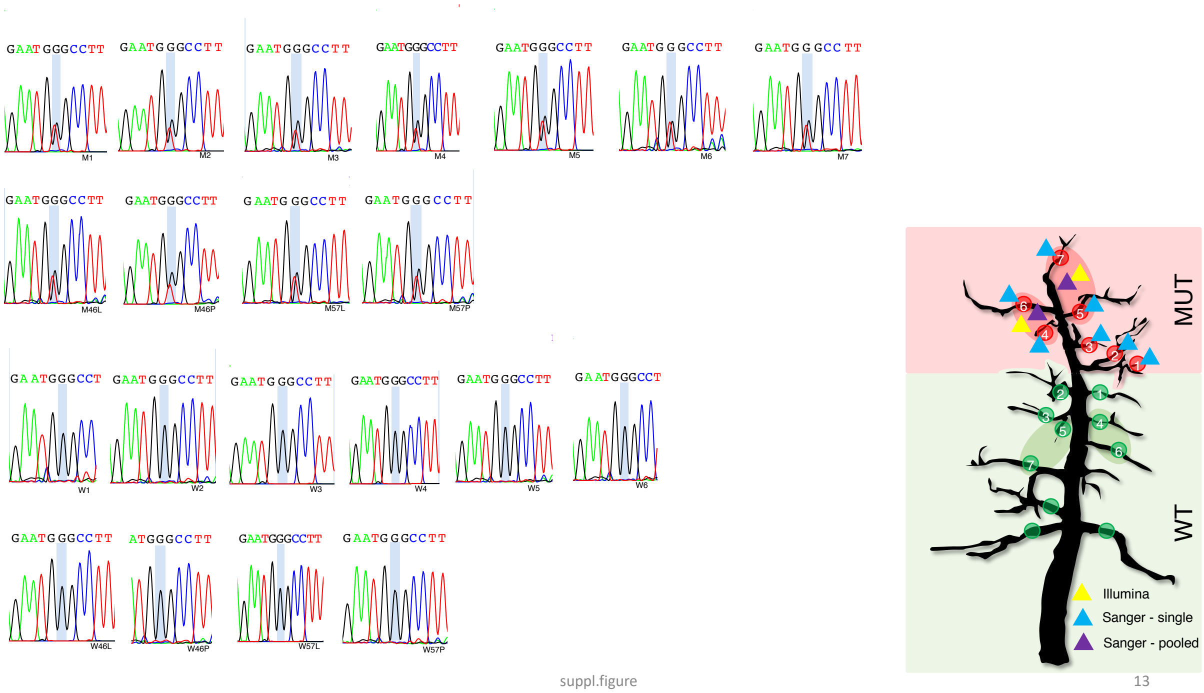


**Supplementary Figure 13. Sanger-seq of mutation No. 46 on fl_3113:G→T (Supplementary Table S2).** MUT: mutant, WT: wildtype. The mutation was found at mutant scaffolds MUT1-7 (as indicated by triangles in blue in the tree sketch), in pooled petiole MUT46P/57P and leaf MUT46L/57L (triangles in purple), while it was not found in any of the individual/pooled wildtype scaffolds. Triangles in yellow indicated the mutations was found by Illumina sequencing of pooled leaf DNA of {4,6} or {5, 7} scaffolds. The mutation site was indicated by the vertical bar in light blue in each Sanger sequencing sample.

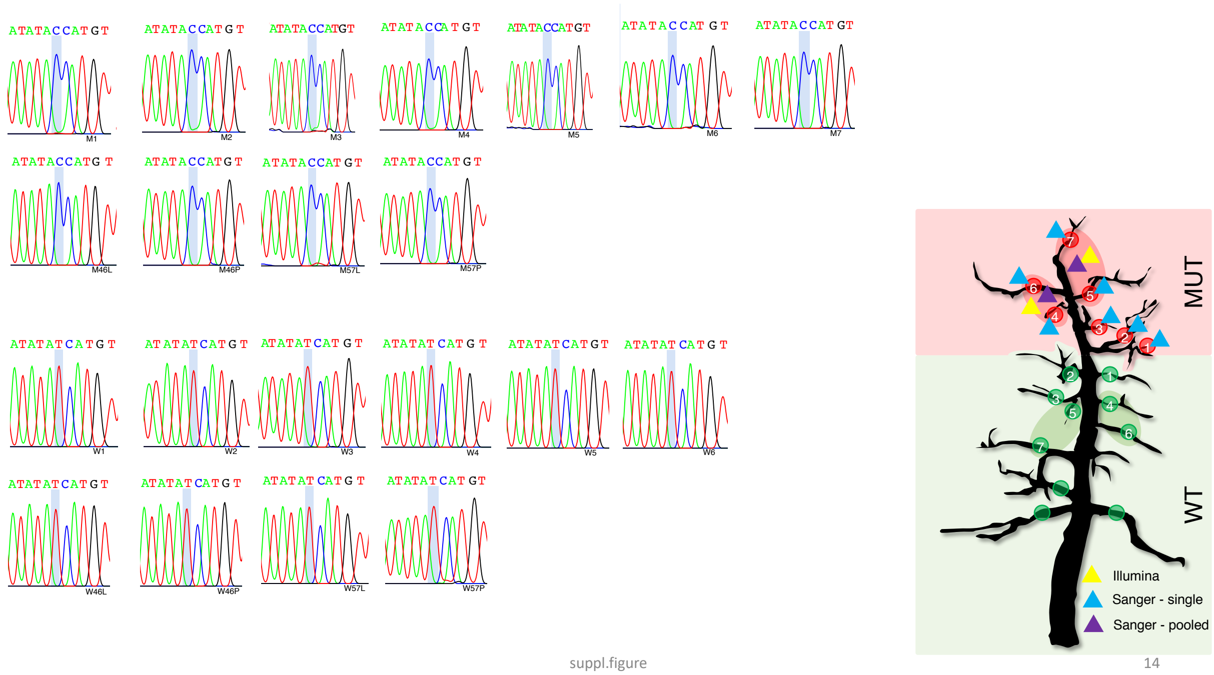


**Supplementary Figure 14. Sanger-seq of mutation No. 47 on fl_3117:T→C (Supplementary Table S2).** MUT: mutant, WT: wildtype. The mutation was found at mutant scaffolds MUT1-7 (as indicated by triangles in blue in the tree sketch), in pooled petiole MUT46P/57P and leaf MUT46L/57L (triangles in purple), while it was not found in any of the individual/pooled wildtype scaffolds. Triangles in yellow indicated the mutations was found by Illumina sequencing of pooled leaf DNA of {4,6} or {5, 7} scaffolds. The mutation site was indicated by the vertical bar in light blue in each Sanger sequencing sample.

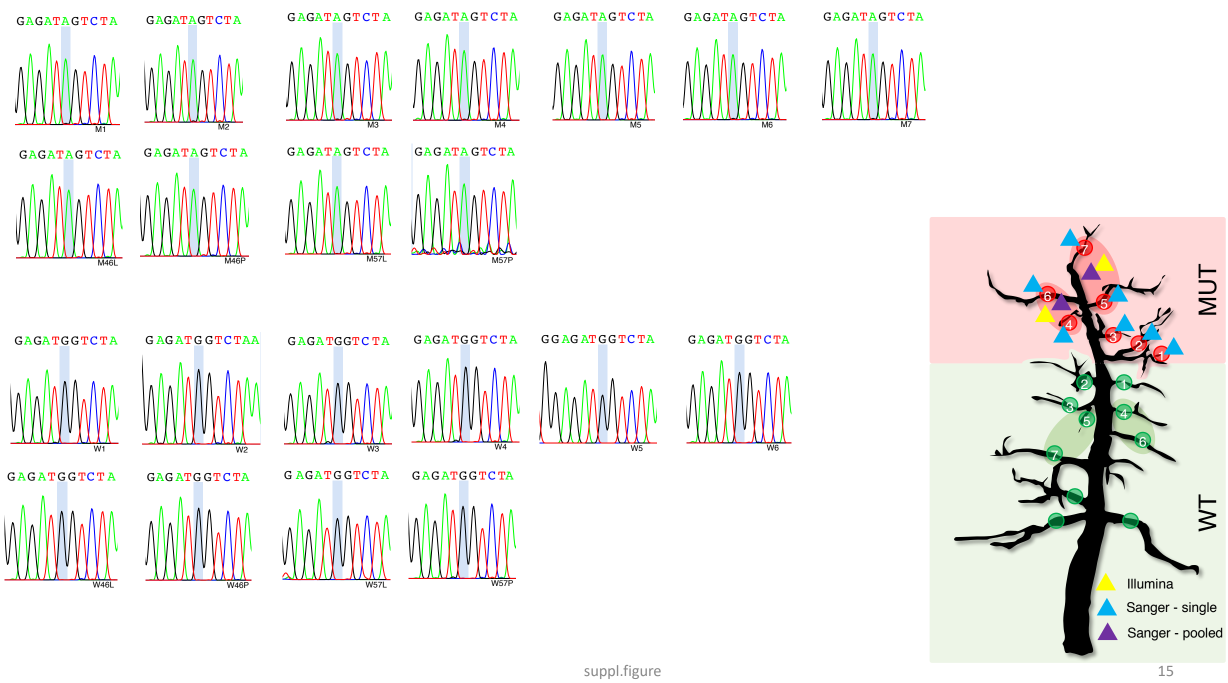


**Supplementary Figure 15. Sanger-seq of mutation No. 50 on fl_3723:G→A (Supplementary Table S2).** MUT: mutant, WT: wildtype. The mutation was found at mutant scaffolds MUT1-7 (as indicated by triangles in blue in the tree sketch), in pooled petiole MUT46P/57P and leaf MUT46L/57L (triangles in purple), while it was not found in any of the individual/pooled wildtype scaffolds. Triangles in yellow indicated the mutations was found by Illumina sequencing of pooled leaf DNA of {4,6} or {5, 7} scaffolds. The mutation site was indicated by the vertical bar in light blue in each Sanger sequencing sample.

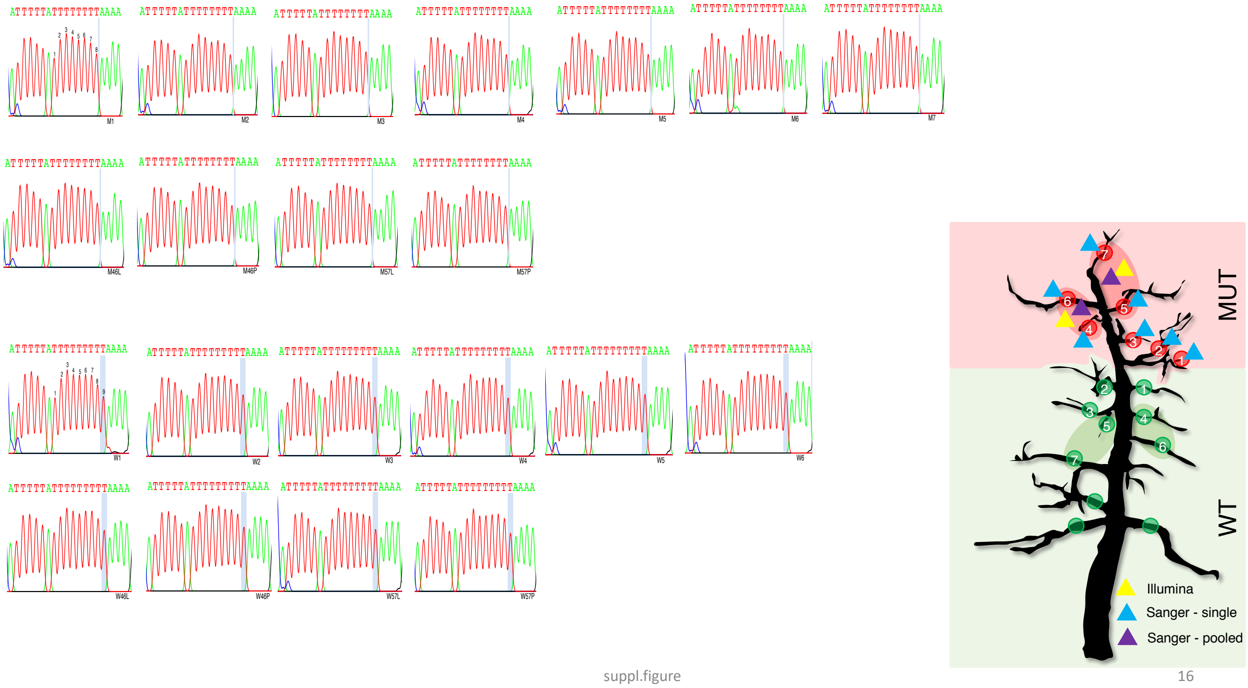


**Supplementary Figure 16. Sanger-seq of mutation No. 53 on fl_37991:T→D (Supplementary Table S2).** MUT: mutant, WT: wildtype. The mutation was found at mutant scaffolds MUT1-7 (as indicated by triangles in blue in the tree sketch), in pooled petiole MUT46P/57P and leaf MUT46L/57L (triangles in purple), while it was not found in any of the individual/pooled wildtype scaffolds. Triangles in yellow indicated the mutations was found by Illumina sequencing of pooled leaf DNA of {4,6} or {5, 7} scaffolds. The mutation site was indicated by the vertical bar in light blue in each Sanger sequencing sample.

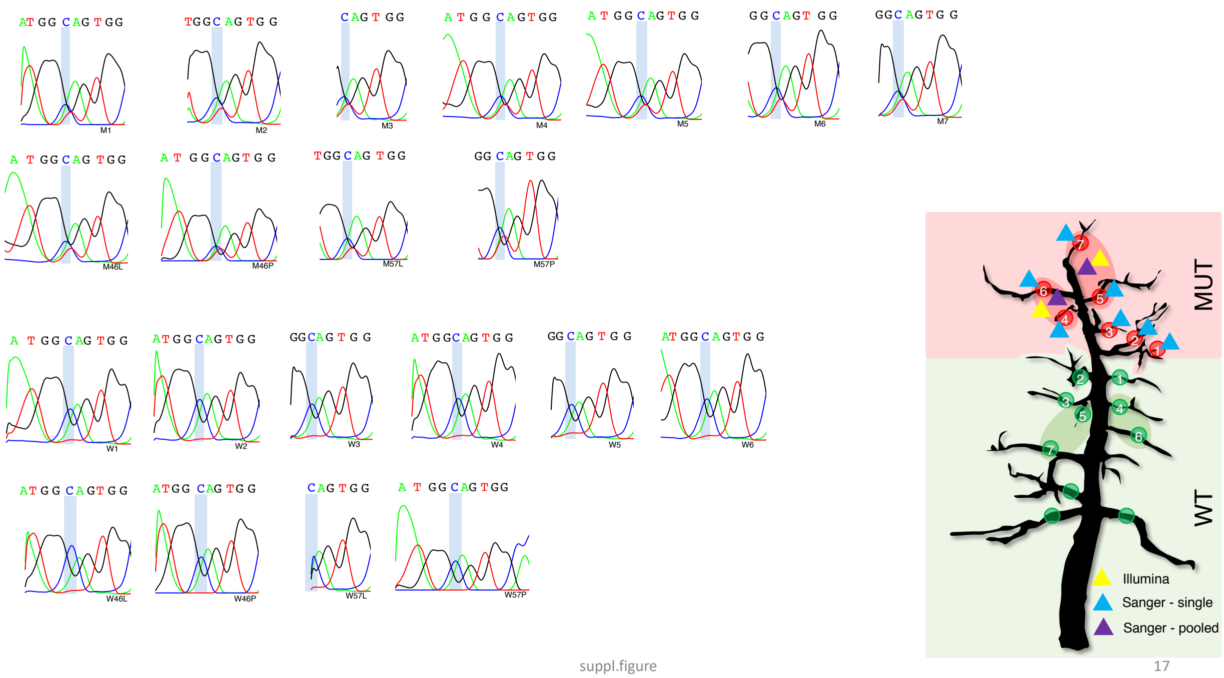


**Supplementary Figure 17. Sanger-seq of mutation No. 93 on fl_44842:C→T (Supplementary Table S2).** MUT: mutant, WT: wildtype. The mutation was found at mutant scaffolds MUT1-7 (as indicated by triangles in blue in the tree sketch), in pooled petiole MUT46P/57P and leaf MUT46L/57L (triangles in purple), while it was not found in any of the individual/pooled wildtype scaffolds. Triangles in yellow indicated the mutations was found by Illumina sequencing of pooled leaf DNA of {4,6} or {5, 7} scaffolds. The mutation site was indicated by the vertical bar in light blue in each Sanger sequencing sample.

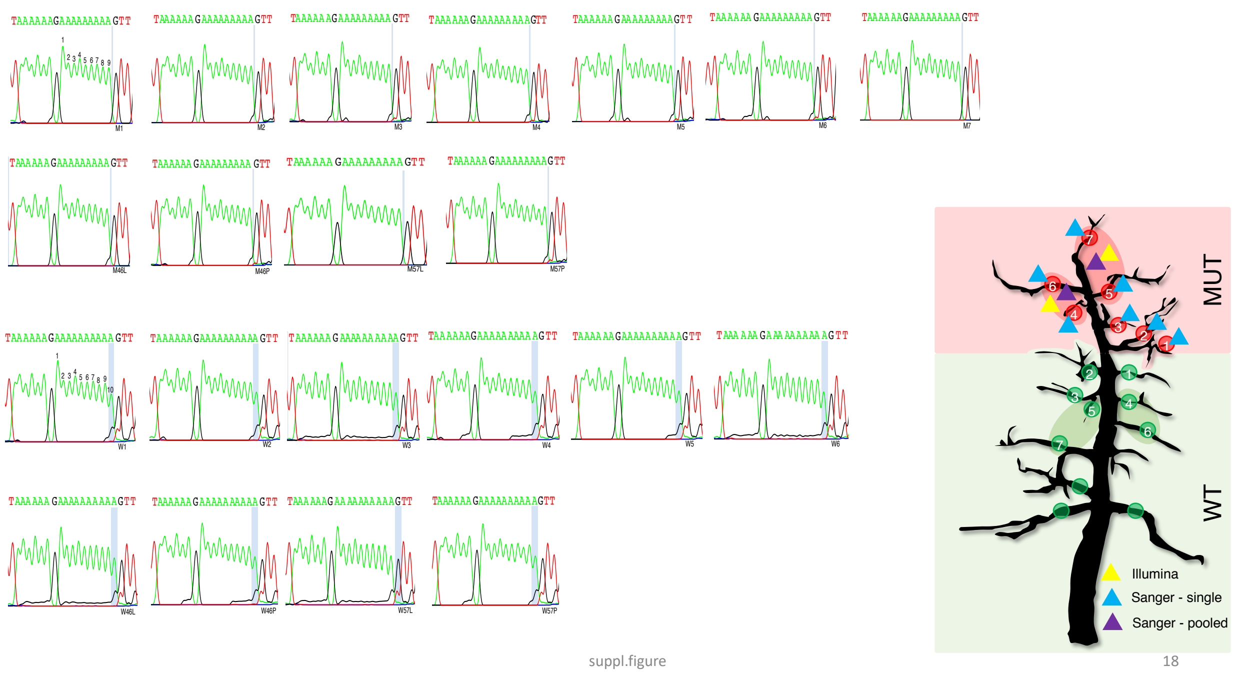


**Supplementary Figure 18. Sanger-seq of mutation No. 76 on fl_66618:A→Deletion (Supplementary Table S2).** MUT: mutant, WT: wildtype. The mutation was found at mutant scaffolds MUT1-7 (as indicated by triangles in blue in the tree sketch), in pooled petiole MUT46P/57P and leaf MUT46L/57L (triangles in purple), while it was not found in any of the individual/pooled wildtype scaffolds. Triangles in yellow indicated the mutations was found by Illumina sequencing of pooled leaf DNA of {4,6} or {5, 7} scaffolds. The mutation site was indicated by the vertical bar in light blue in each Sanger sequencing sample.

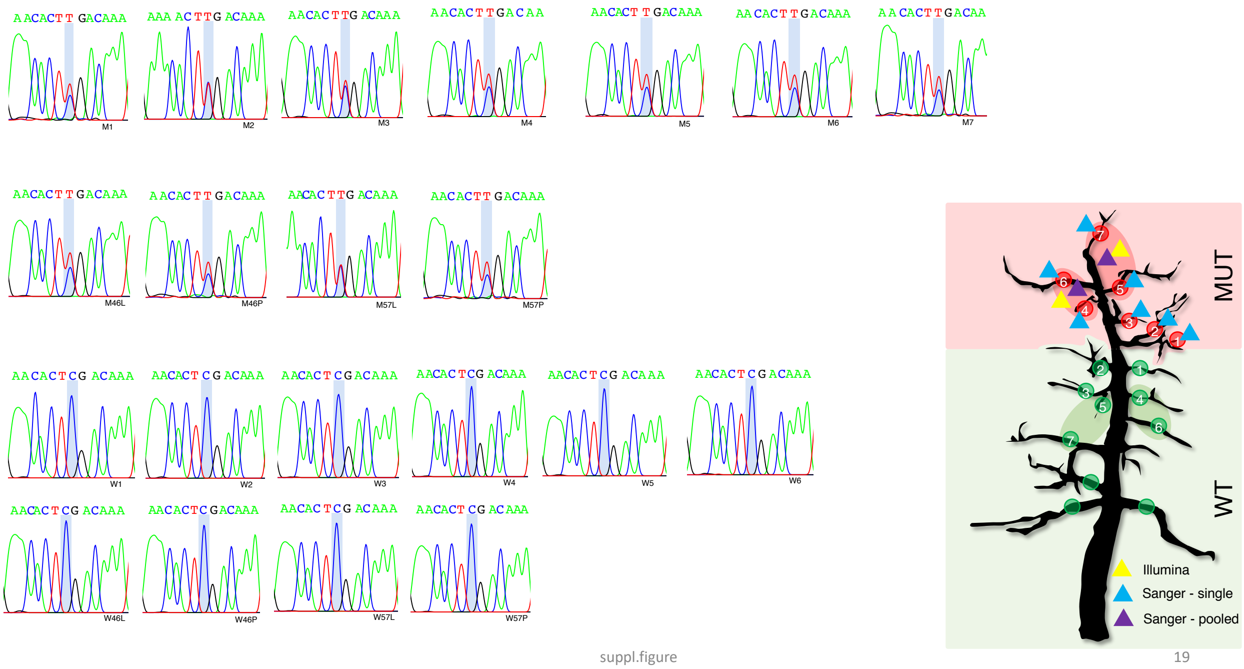


**Supplementary Figure 19. Sanger-seq of mutation No. 80 on fl_73045:C→T (Supplementary Table S2).** MUT: mutant, WT: wildtype. The mutation was found at mutant scaffolds MUT1-7 (as indicated by triangles in blue in the tree sketch), in pooled petiole MUT46P/57P and leaf MUT46L/57L (triangles in purple), while it was not found in any of the individual/pooled wildtype scaffolds. Triangles in yellow indicated the mutations was found by Illumina sequencing of pooled leaf DNA of {4,6} or {5, 7} scaffolds. The mutation site was indicated by the vertical bar in light blue in each Sanger sequencing sample.

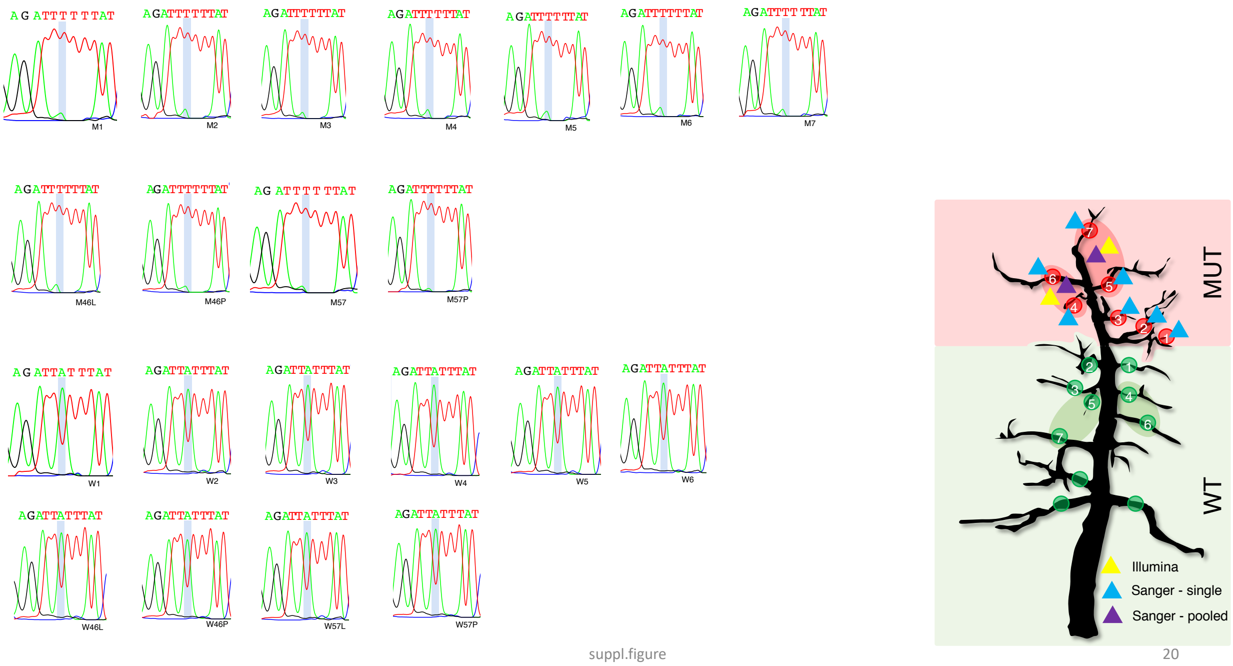


**Supplementary Figure 20. Sanger-seq of mutation No. 91 on fl_8965:A→T (Supplementary Table S2).** MUT: mutant, WT: wildtype. The mutation was found at mutant scaffolds MUT1-7 (as indicated by triangles in blue in the tree sketch), in pooled petiole MUT46P/57P and leaf MUT46L/57L (triangles in purple), while it was not found in any of the individual/pooled wildtype scaffolds. Triangles in yellow indicated the mutations was found by Illumina sequencing of pooled leaf DNA of {4,6} or {5, 7} scaffolds. The mutation site was indicated by the vertical bar in light blue in each Sanger sequencing sample.

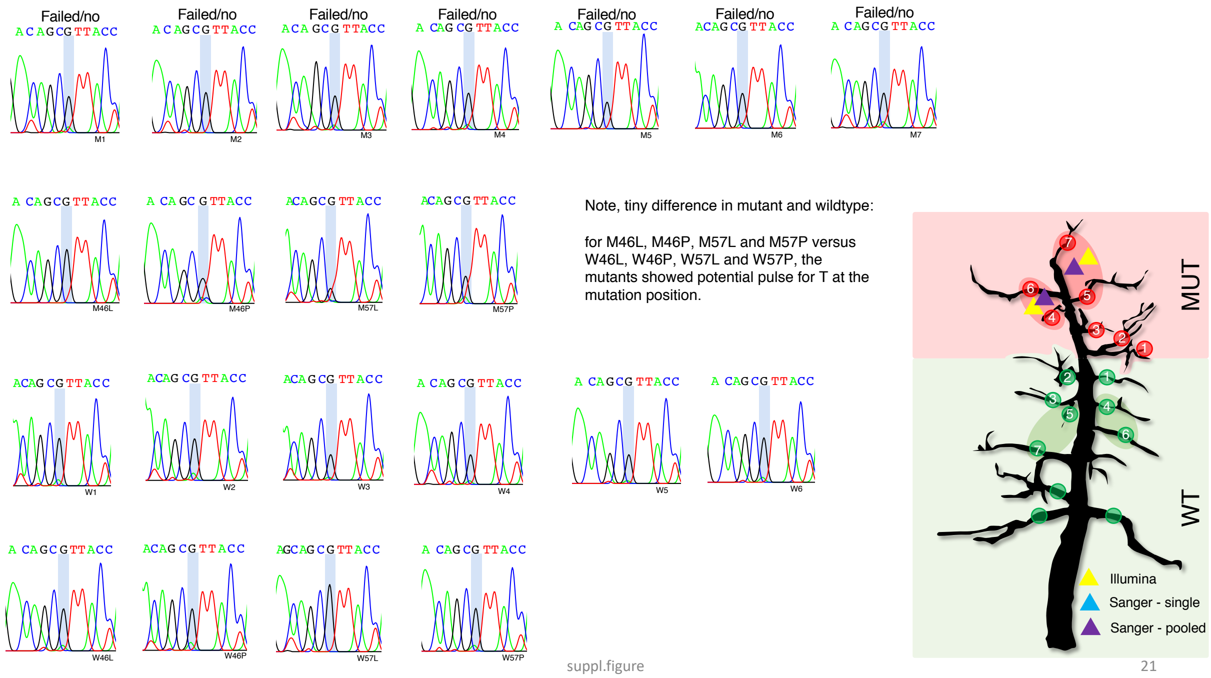


**Supplementary Figure 21. Sanger-seq of mutation No. 56 on fl_44093:G→T (Supplementary Table S2).** MUT: mutant, WT: wildtype. The mutation was only found in mutant samples of pooled petiole MUT46P/57P and leaf MUT46L/57L (triangles in purple), while it was not found in any of the individual/pooled wildtype scaffolds. Triangles in yellow indicated the mutations was found by Illumina sequencing of pooled leaf DNA of {4,6} or {5, 7} scaffolds. The mutation site was indicated by the vertical bar in light blue in each Sanger sequencing sample. "Failed/no" indicated cases where Sanger sequencing did not work or no alternative allele could be found.

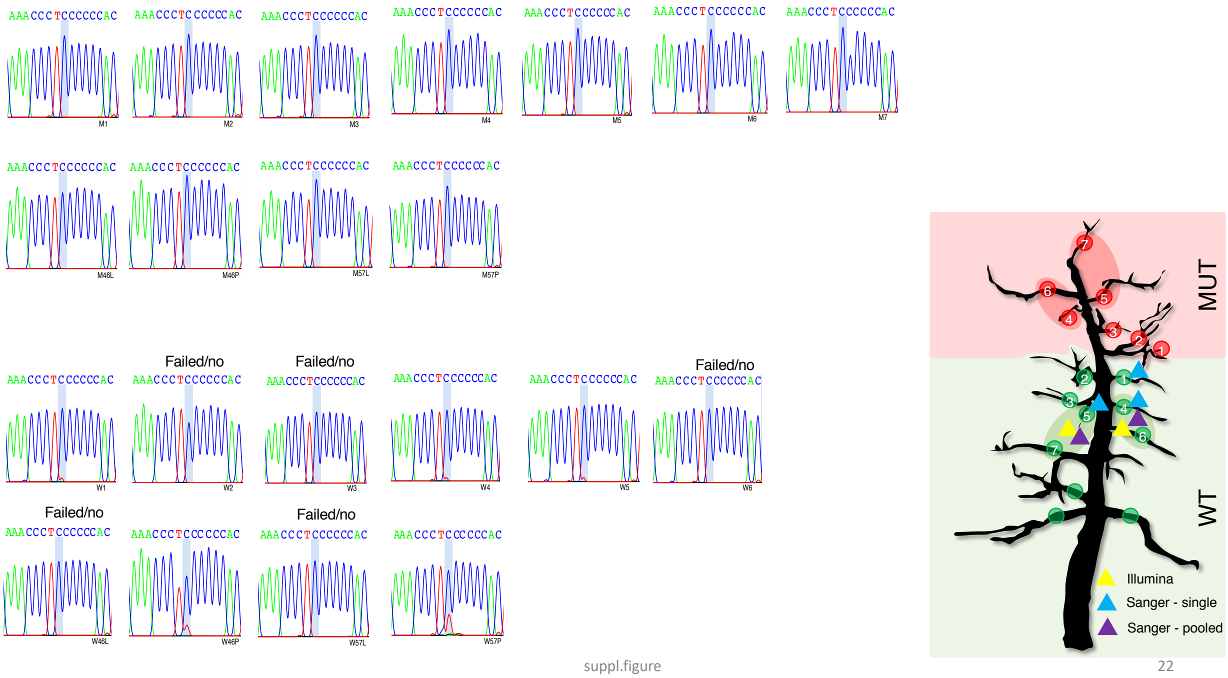


**Supplementary Figure 22. Sanger-seq of mutation No. 32 on fl_20587:T→C (Supplementary Table S2).** MUT: mutant, WT: wildtype. The mutation was found at wildtype scaffolds WT1,4,5 (as indicated by triangles in blue in the tree sketch), in pooled petiole WT46P/57P (triangles in purple), while it was not found in any of the individual/pooled mutant scaffolds. Triangles in yellow indicated the mutations was found by Illumina sequencing of pooled leaf DNA of {4,6} or {5, 7} scaffolds. The mutation site was indicated by the vertical bar in light blue in each Sanger sequencing sample. “Failed/no” indicated cases where Sanger sequencing did not work or no alternative allele could be found.

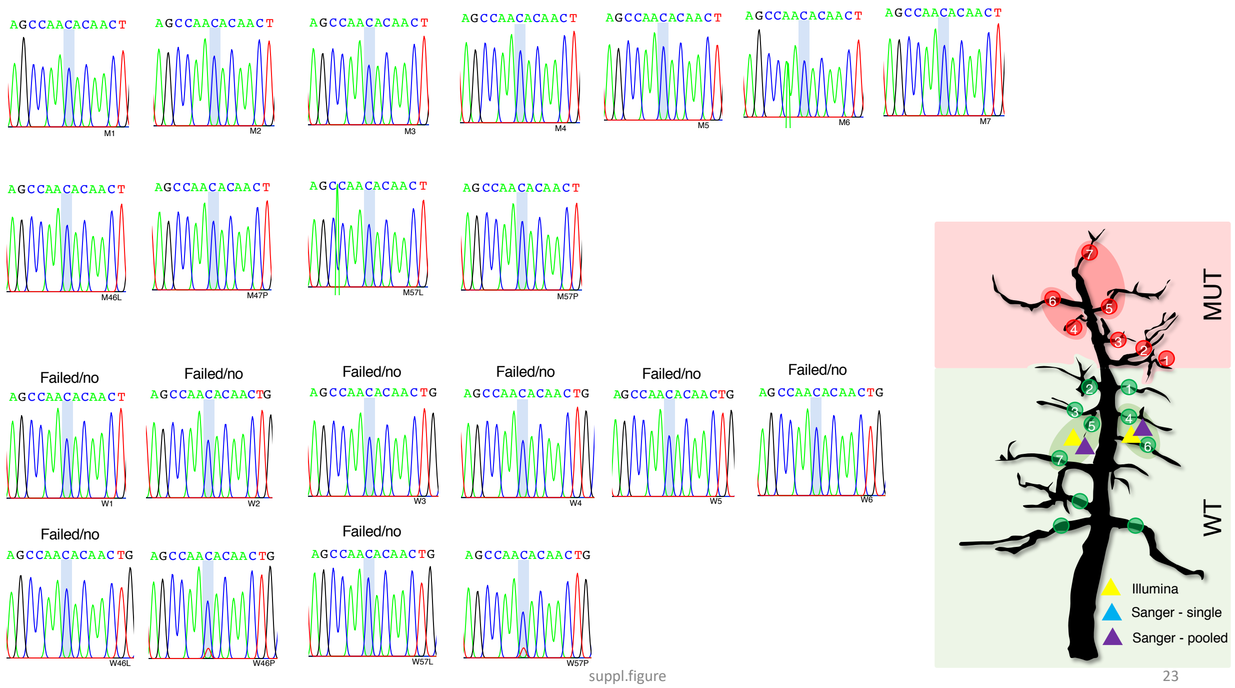


**Supplementary Figure 23. Sanger-seq of mutation No. 45 on fl_3018:T→C (Supplementary Table S2).** MUT: mutant, WT: wildtype. The mutation was found in pooled petiole Wt46P/57P (triangles in purple), while it was not found in any of the individual/pooled mutant scaffolds. Triangles in yellow indicated the mutations was found by Illumina sequencing of pooled leaf DNA of {4,6} or {5, 7} scaffolds. The mutation site was indicated by the vertical bar in light blue in each Sanger sequencing sample. “Failed/no” indicated cases where Sanger sequencing did not work or no alternative allele could be found.

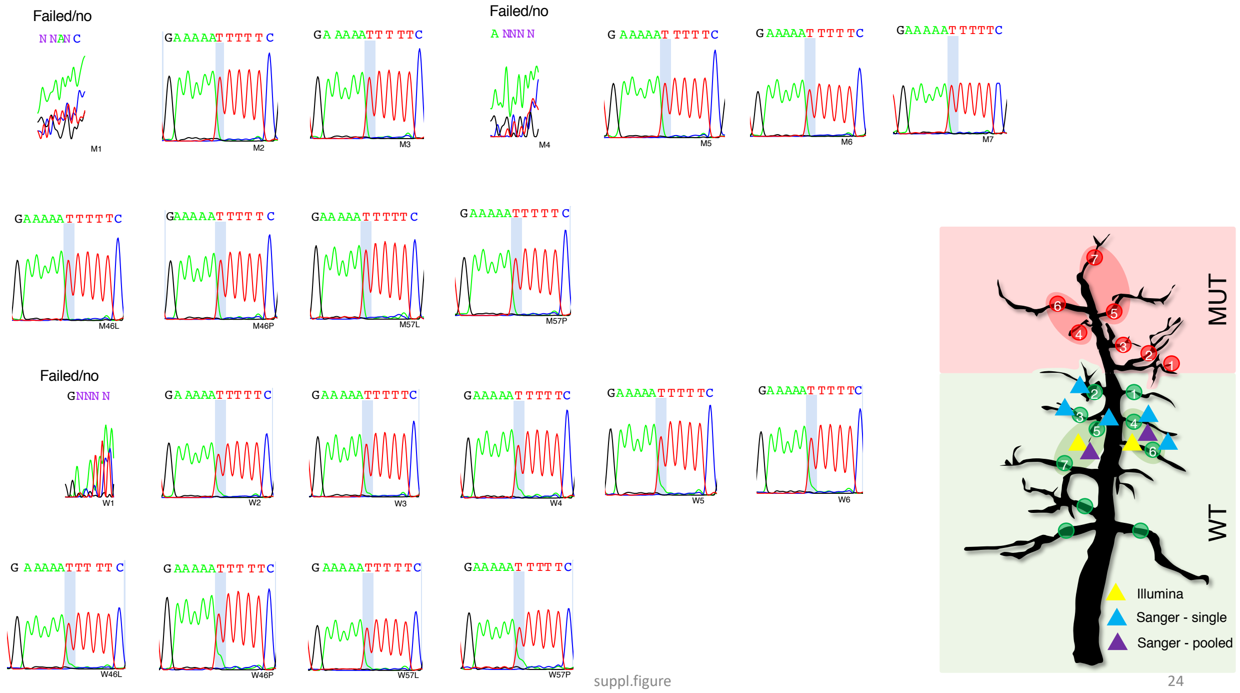


**Supplementary Figure 24. Sanger-seq of mutation No. 73 on fl_6517:A→T (Supplementary Table S2).** MUT: mutant, WT: wildtype. The mutation was found at wildtype scaffolds WT2-6 (as indicated by triangles in blue in the tree sketch), in pooled petiole WT46P/57P and WT46L/57L (triangles in purple), while it was not found in any of the individual/pooled mutant scaffolds. Triangles in yellow indicated the mutations was found by Illumina sequencing of pooled leaf DNA of {4,6} or {5, 7} scaffolds. The mutation site was indicated by the vertical bar in light blue in each Sanger sequencing sample. "Failed/no" indicated cases where Sanger sequencing did not work or no alternative allele could be found.

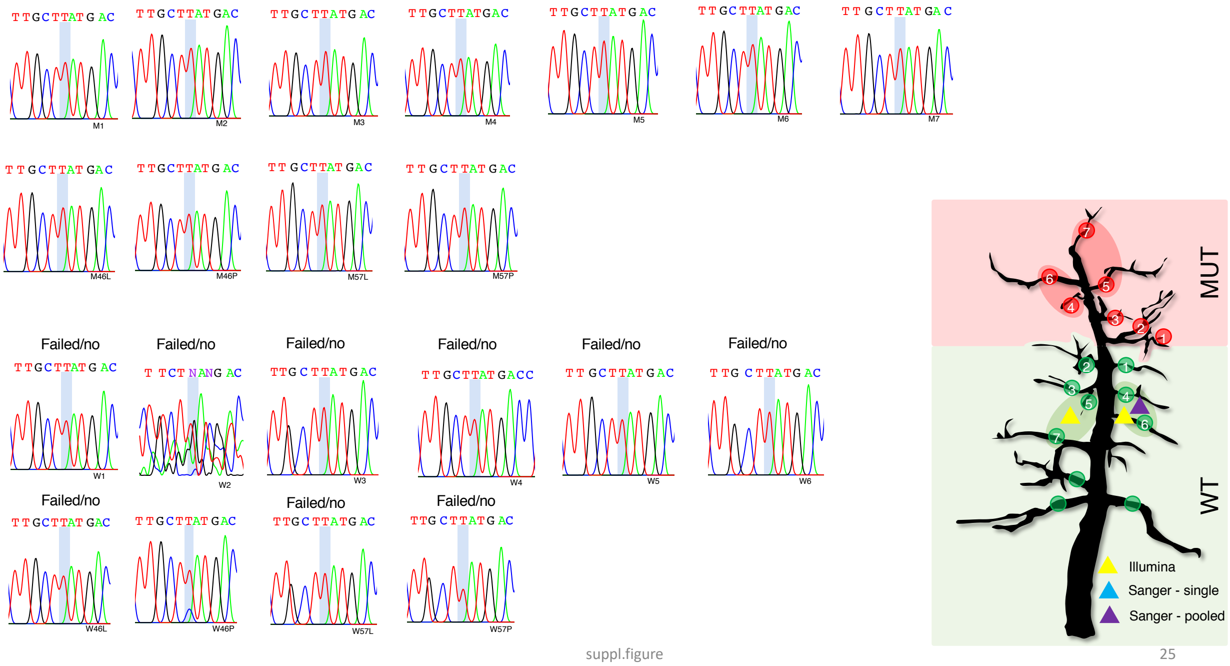


**Supplementary Figure 25. Sanger-seq of mutation No. 77 on fl_67102 (Supplementary Table S2).** MUT: mutant, WT: wildtype. The mutation was only found in pooled petiole WT46P (triangles in purple), while it was not found in any of the individual/pooled mutant scaffolds. Triangles in yellow indicated the mutations was found by Illumina sequencing of pooled leaf DNA of {4,6} or {5, 7} scaffolds. The mutation site was indicated by the vertical bar in light blue in each Sanger sequencing sample. “Failed/no” indicated cases where Sanger sequencing did not work or no alternative allele could be found.

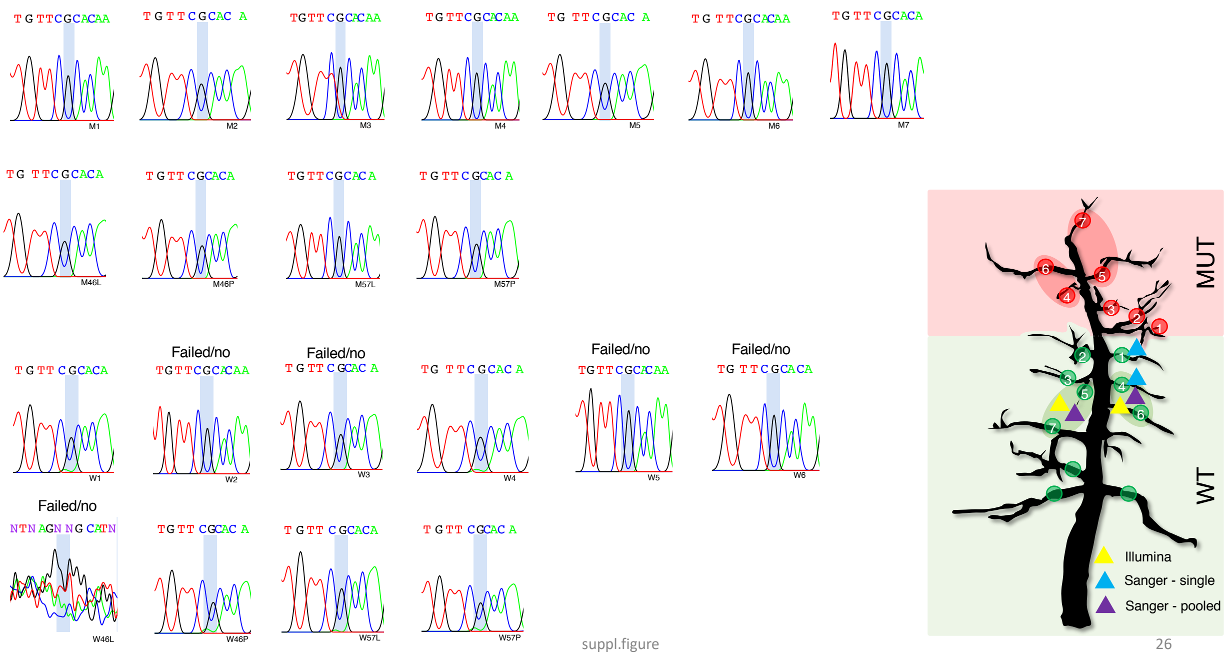


**Supplementary Figure 26. Sanger-seq of mutation No. 81 on fl_74:A→G (Supplementary Table S2).** MUT: mutant, WT: wildtype. The mutation was found at wildtype scaffolds WT1,4 (as indicated by triangles in blue in the tree sketch), in pooled petiole WT46P/57P and WT57L (triangles in purple), while it was not found in any of the individual/pooled mutant scaffolds. Triangles in yellow indicated the mutations was found by Illumina sequencing of pooled leaf DNA of {4,6} or {5, 7} scaffolds. The mutation site was indicated by the vertical bar in light blue in each Sanger sequencing sample. "Failed/no" indicated cases where Sanger sequencing did not work or no alternative allele could be found.

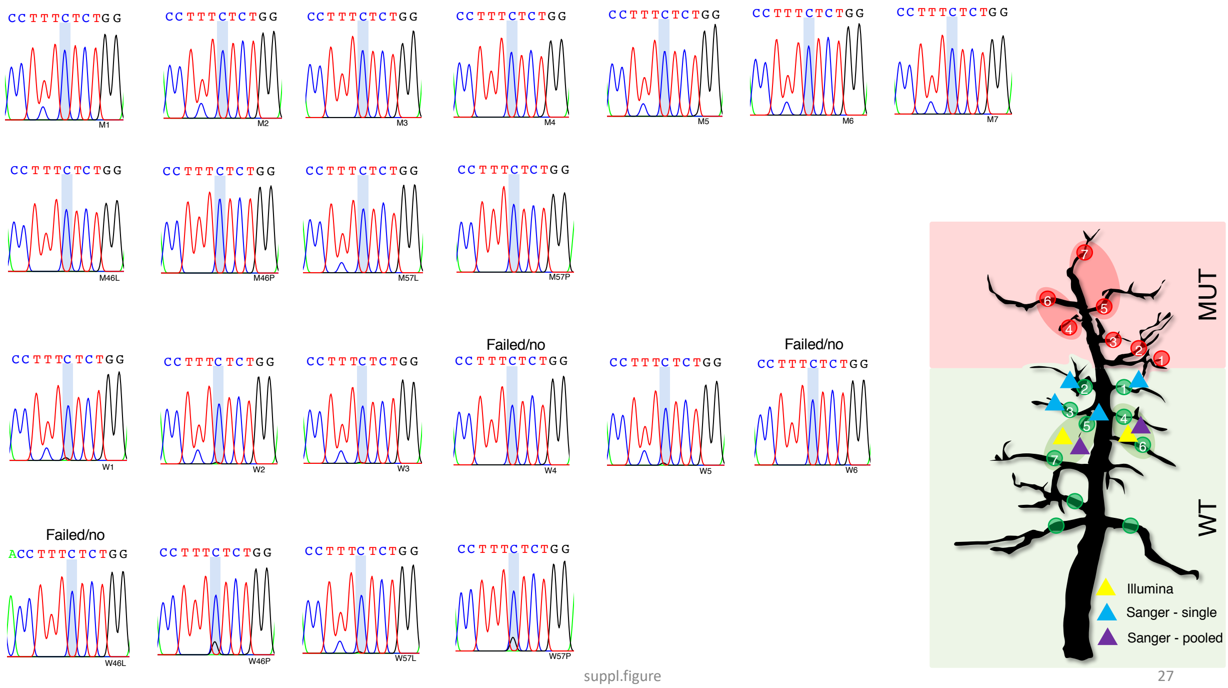


**Supplementary Figure 27. Sanger-seq of mutation No. 43 on fl_2910:G→C (Supplementary Table S2).** MUT: mutant, WT: wildtype. The mutation was found at wildtype scaffolds WT1,2,3,5 (as indicated by triangles in blue in the tree sketch), in pooled petiole WT46P/57P and pooled leaf WT57L (triangles in purple), while it was not found in any of the individual/pooled mutant scaffolds. Triangles in yellow indicated the mutations was found by Illumina sequencing of pooled leaf DNA of {4,6} or {5, 7} scaffolds. The mutation site was indicated by the vertical bar in light blue in each Sanger sequencing sample. “Failed/no” indicated cases where Sanger sequencing did not work or no alternative allele could be found.

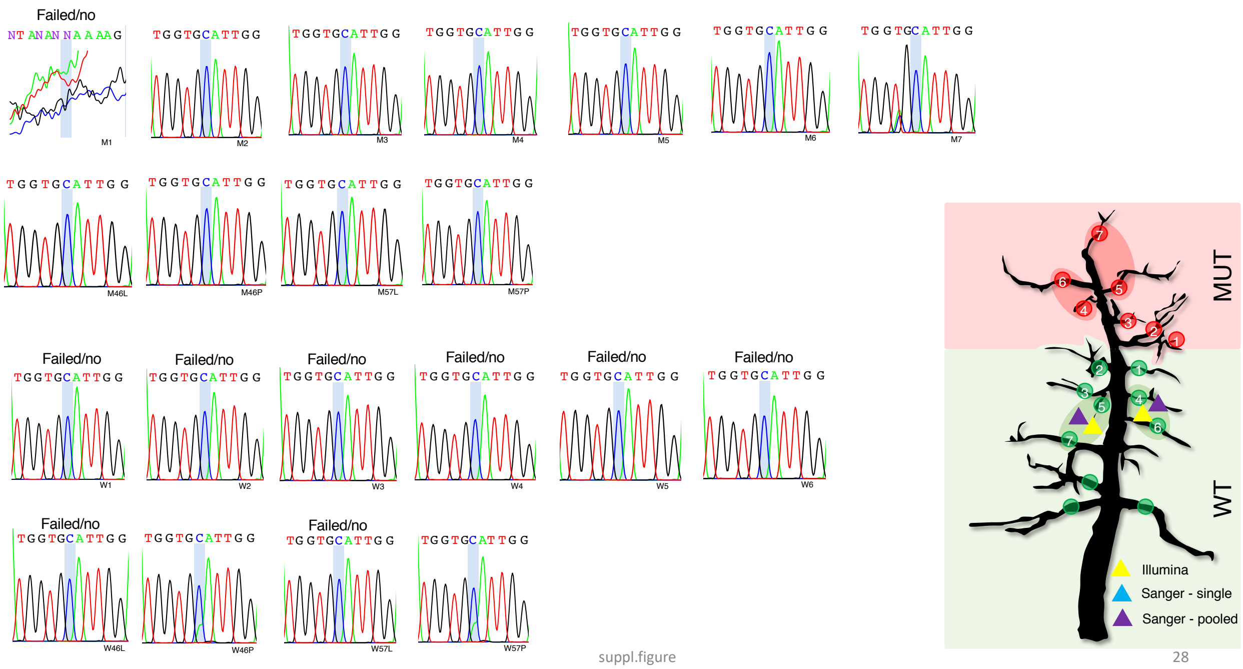


**Supplementary Figure 28. Sanger-seq of mutation No. 62 on fl_4941:A→C (Supplementary Table S2).** MUT: mutant, WT: wildtype. The mutation was only found in pooled petiole WT46P/57P (triangles in purple), while it was not found in any of the individual/pooled mutant scaffolds. Triangles in yellow indicated the mutations was found by Illumina sequencing of pooled leaf DNA of {4,6} or {5, 7} scaffolds. The mutation site was indicated by the vertical bar in light blue in each Sanger sequencing sample. “Failed/no” indicated cases where Sanger sequencing did not work or no alternative allele could be found.

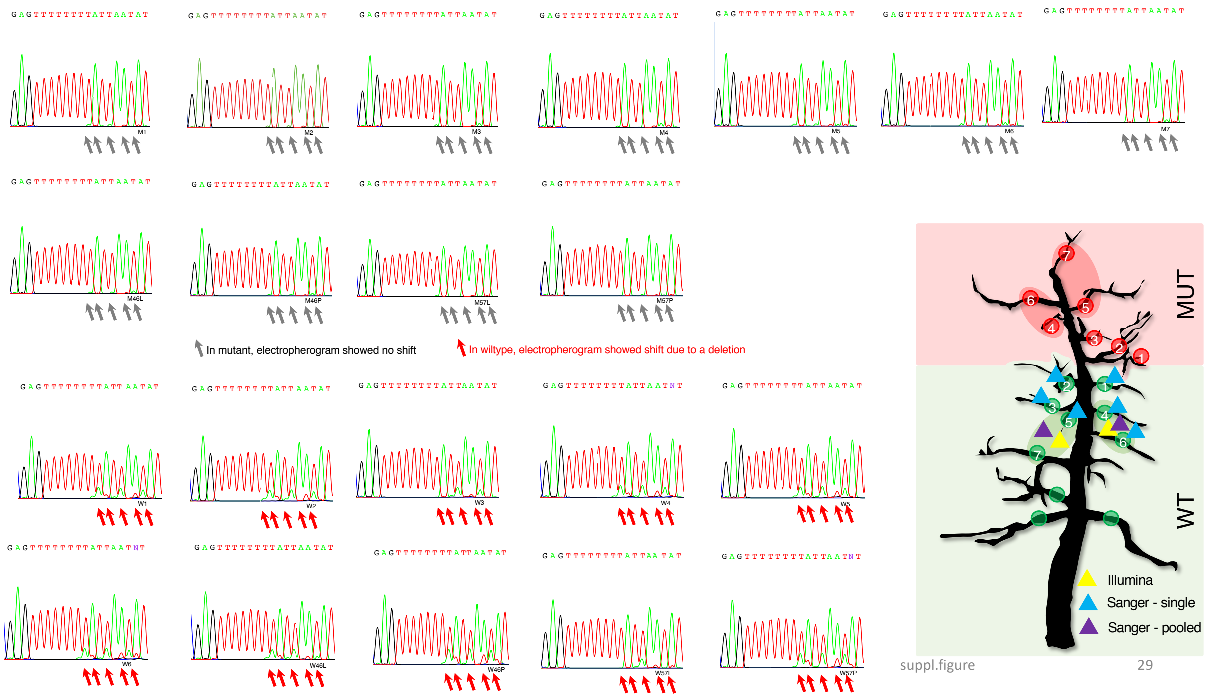


**Supplementary Figure 29. Sanger-seq of mutation No. 65 on fl_5351:Deletion→A (Supplementary Table S2).** MUT: mutant, WT: wildtype. The mutation was found at wildtype scaffolds WT1-6 (as indicated by triangles in blue in the tree sketch), in pooled petiole WT46P/57P and WT46L/57L (triangles in purple), while it was not found in any of the individual/pooled mutant scaffolds. Triangles in yellow indicated the mutations was found by Illumina sequencing of pooled leaf DNA of {4,6} or {5, 7} scaffolds. The mutation site was indicated by the vertical bar in light blue in each Sanger sequencing sample. “Failed/no” indicated cases where Sanger sequencing did not work or no alternative allele could be found.

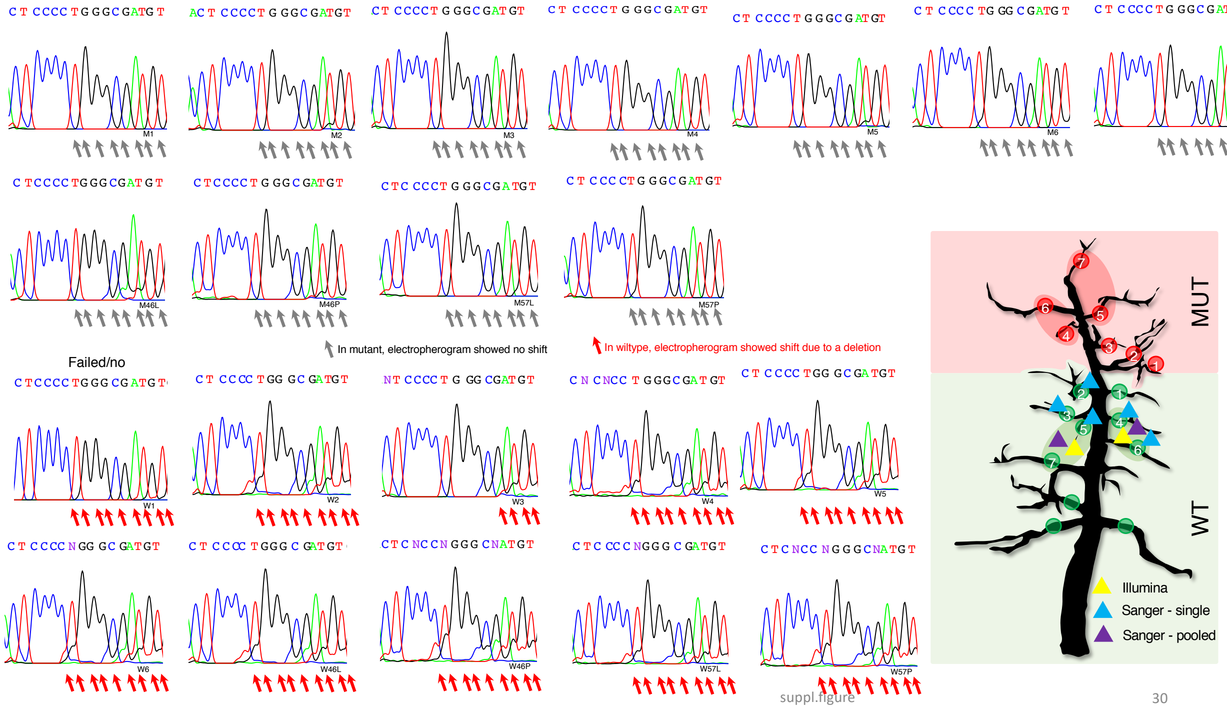


**Supplementary Figure 30. Sanger-seq of mutation No. 88 on fl_86445:Deletion→C (Supplementary Table S2).** MUT: mutant, WT: wildtype. The mutation was found at wildtype scaffolds WT2-6 (as indicated by triangles in blue in the tree sketch), in pooled petiole WT46P/57P and WT46L/57L (triangles in purple), while it was not found in any of the individual/pooled mutant scaffolds. Triangles in yellow indicated the mutations was found by Illumina sequencing of pooled leaf DNA of {4,6} or {5, 7} scaffolds. The mutation site was indicated by the vertical bar in light blue in each Sanger sequencing sample. “Failed/no” indicated cases where Sanger sequencing did not work or no alternative allele could be found.

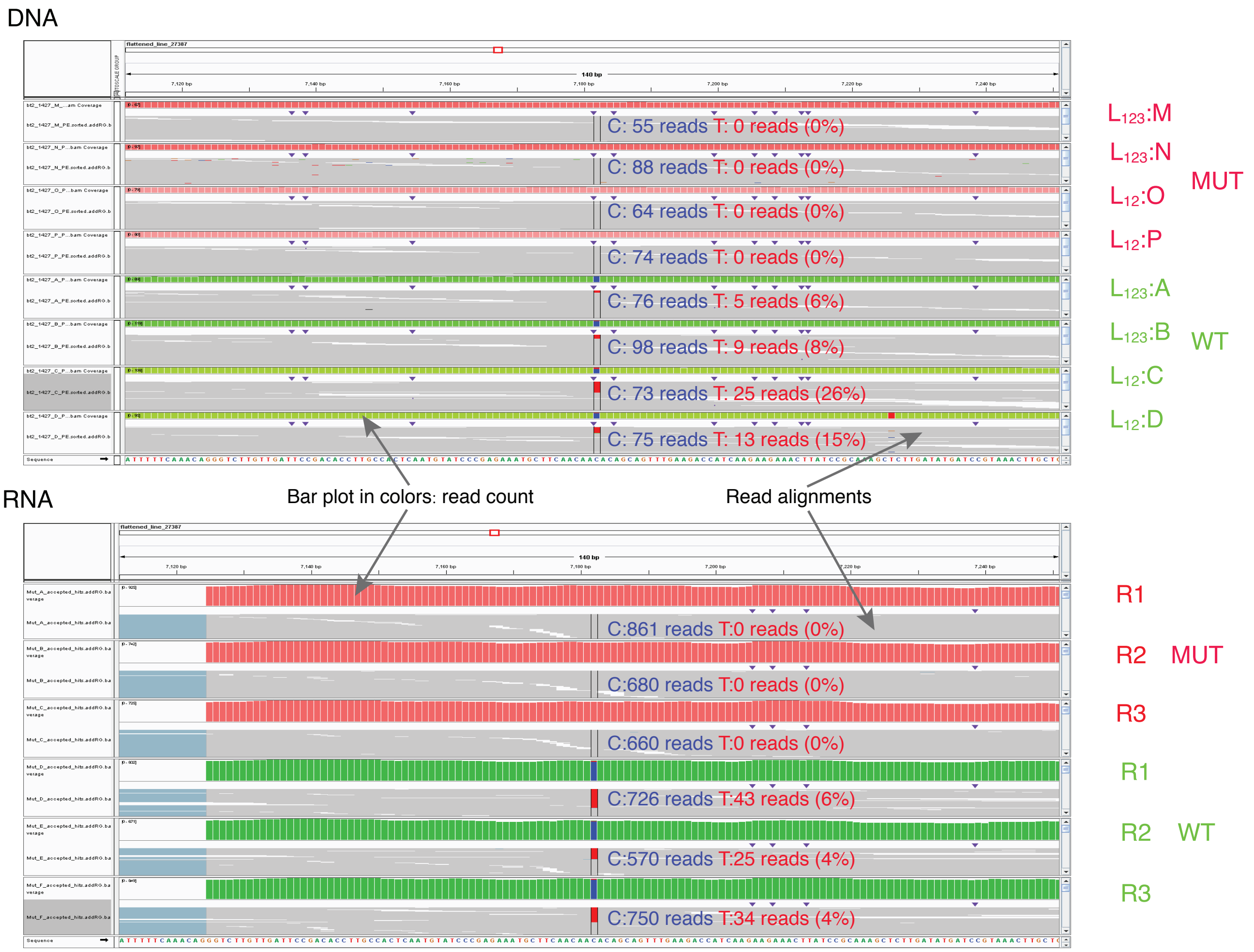


**Supplementary Figure 31. Validation of the mutation No. 39 at fl_27387:7182 hitting the HSP90 gene in *RubyMac*.** MUT: mutant, WT: wildtype. According to the Illumina based sequencing of leaf DNA (top panel), in wildtype samples (WT A-D in green), there were 6-8% reads sequenced from $L_{123}$ samples versus 15-26% reads sequenced from $L_{12}$ samples showing the alternative allele, while no such reads observed in mutant samples (MUT M-P in red). This suggested the alternative allele was only carried by cells in $L_1$ or $L_2$, which could result in low allele frequency in $L_{123}$ DNA. This low-frequency mutation was not detected by Sanger sequencing. However, in wildtype replicates (WT R1-3 in green of bottom panel), there were 4-6% (or in absolute numbers of 25-43) of the RNA-seq reads of HSP90 that carried the mutation, while not a single such read was found in mutant replicates (MUT R1-3 in red), suggesting this was a real difference between wildtype and mutant samples.
